# Two intrinsic timing mechanisms set start and end times for dendritic arborization of a nociceptive neuron

**DOI:** 10.1101/2021.08.31.458402

**Authors:** Nobuko Suzuki, Yan Zou, HaoSheng Sun, Kelsie Eichel, Meiyu Shao, Kang Shen, Chieh Chang

**Affiliations:** Department of Biological Sciences, University of Illinois at Chicago, Chicago, IL 60607, USA; School of Life Science and Technology, ShanghaiTech University, Shanghai 201210, China; Department of Cell, Developmental and Integrative Biology, University of Alabama at Birmingham, Birmingham, AL 35233, USA; Howard Hughes Medical Institute, Department of Biology, Stanford University, Stanford, CA 94305, USA

**Keywords:** the *lin-4-lin-14* pathway, the *lin-28-let-7-lin-41* pathway, developmental timing genes, heterochronic genes, neuronal timers, temporal regulation, dendrite arborization, PVD neurons, *Caenorhabditis elegans*

## Abstract

Choreographic dendritic arborization takes place within a defined time frame, but the timing mechanism is currently not known. Here, we report that the precisely timed *lin-4-lin-14* regulatory circuit triggers an initial dendritic growth activity whereas the precisely timed *lin-28-let-7-lin-41* regulatory circuit signals a subsequent developmental decline in dendritic growth ability, hence restricting dendritic arborization within a set time frame. Loss-of-function mutations in the *lin-4* microRNA gene cause limited dendritic outgrowth whereas loss-of-function mutations in its direct target, the *lin-14* transcription factor gene, cause precocious and excessive outgrowth. In contrast, loss-of-function mutations in the *let-7* microRNA gene prevent a developmental decline in dendritic growth ability whereas loss-of-function mutations in its upstream regulator, the *lin-28* RNA-binding protein gene, and its direct target, the *lin-41* tripartite motif protein gene, cause further decline. *lin-4* and *let-7* regulatory circuits are expressed in the right place at the right time to set start and end times for PVD dendritic arborization. Replacing the endogenous *lin-4* promoter at the *lin-4* locus with a late-onset *let-7* promoter delays PVD dendrite arborization whereas replacing the endogenous *let-7* promoter at the *let-7* locus with an early-onset *lin-4* promoter causes precocious decline in dendritic growth ability in PVD neurons. Our results indicate that the *lin-4-lin-14* and the *lin-28-let-7-lin-41* regulatory circuits control the timing of dendrite arborization through antagonistic regulation of the DMA-1 receptor level on dendrites. The LIN-14 transcription factor directly represses the *dma-1* gene expression through a transcriptional means whereas the LIN-41 tripartite motif protein indirectly activates the *dma-1* gene expression through a post-transcriptional means.

## Introduction

Studies on temporal control of cell cycle progression, circadian rhythm, and segmentation have frequently converged on the concept of biological oscillators (Bargiello et al., 1984; Cooke and Zeeman, 1976; Hardin et al., 1990; Horikawa et al., 2006; Kondo, 2007; Liu et al., 1992; Morgan, 1997; Nakajima et al., 2005; Oates et al., 2012; Price et al., 1998; Reppert and Weaver, 2002; Siwicki et al., 1988; Vosshall et al., 1994; Zehring et al., 1984). Biological oscillators are systems of molecules with various levels of expression and activity that act as molecular “clocks” that determine biological rhythms resilient to changes in external environments. Understanding these molecular oscillators gives a glimpse into the mechanism of timing control for the cyclical nature of these processes. However, timing control for non-cyclical biological processes is less understood. Most non-cyclical events in non-neuronal cells are transient, which makes it difficult to study their temporal regulation. In contrast, differentiation of neurons into complex structures occurs on longer timescales, thereby giving us a window to peek into non-cyclical timing mechanisms.

Although abundant knowledge has been learnt in past decades on temporal control of cell fate specification of neurons (Faunes and Larraín, 2016; Pereira et al., 2019; Rougvie and Moss, 2013), less is known for timing of their wiring to give rise to complex neuronal circuits and for timing of their plasticity. In the nervous system, establishment of neuronal connectivity during development and decline in neuronal plasticity during aging are controlled with temporal precision, but the timing mechanisms are largely unknown until recently. The heterochronic pathways are important temporal regulators of animal development and involve a number of microRNA-regulated post-transcriptional genetic circuits, including important interactions between the *lin-4* microRNA and its direct target, the *lin-14* transcription factor gene (Lee et al., 1993; Wightman et al., 1993) and between the *let-7* microRNA and its direct target, the *lin-41* tripartite motif (TRIM) protein gene (Pasquinelli et al., 2000; Slack et al., 2000). Since the discovery of the *lin-4-lin-14* and the *let-7-lin-41* regulatory circuits broadly expressed in the nervous system (Shih and Chang, 2021; Zou et al., 2012; Zou et al., 2013), evidences indicating a widespread role of the *lin-4-lin-14* and the *let-7-lin-41* regulatory circuits in timing neuronal assembly and plasticity start to accumulate. Distinct microRNA regulatory circuits were shown to control orderly neuronal connectivity and regeneration potential decline (Chiu et al., 2014; Chiu and Chang, 2013; Zou et al., 2012; Zou et al., 2013). The *lin-4* microRNA and the LIN-14 transcription factor regulate transition of sequential events in AVM neuronal connectivity. Up-regulation of *lin-4* and down-regulation of *lin-14* signal the end of netrin-mediated axon pathfinding to allow synapse formation in AVM neurons (Chiu et al., 2014; Zou et al., 2012). Similar functions of *lin-4* and *lin-14* in temporal control of axon pathfinding have been reported in HSN and PLM neurons (Olsson-Carter and Slack, 2010; Xu and Quinn, 2016). The *lin-4-lin-14* regulatory circuit also temporally controls synaptic rewiring of postmitotic DD motor neurons (Hallam and Jin, 1998; Howell et al., 2015). Furthermore, in PVT interneurons, *lin-14* temporally regulates onset of *zig* gene expression, which is required for maintenance of ventral nerve cord structure (Aurelio et al., 2003). The *let-7* microRNA and the LIN-41 tripartite motif protein control the timing of a post-differentiation event in AVM neurons (Chiu and Chang, 2013; Zou et al., 2013). The progressive increase of *let-7* and the progressive decrease of *lin-41* contribute to a normal developmental decline in AVM axon regeneration. Collectively, *lin-4* and *let-7* microRNAs have been recently shown to regulate the establishment of axodendritic polarity in the DA9 motor neuron early in development and its maintenance at the adult stage, respectively (Armakola and Ruvkun, 2019). Despite advances in understanding the temporal transition of neuronal connectivity and regeneration decline, intrinsic timing mechanisms that control choreographic dendritic arborization, an integral part of establishing a functional neural circuit, are still elusive.

Dendritic arborization of the PVD nociceptive neurons takes place within a defined time period starting at the L2 stage and ending at the young adult stage during *C. elegans* development. The PVD dendritic arbor is established by a complex but well-ordered array of non-overlapping sister dendrites. The creation of this structure involves a sequential series of branching decisions, which makes PVD neurons an ideal model system to study timing mechanisms of dendritic arborization. A 1° dendrite extends from the PVD cell body along the A/P axis at the location of the lateral nerve fascicle. Orthogonal arrays of 2°, 3°, and 4° dendritic branches envelop the animal in a manner that alternates between the D/V and the A/P axes to produce an elaborate network of sensory processes (Inberg et al., 2019; Sundararajan et al., 2019). The well-ordered dendrite branches are likened to menorahs, branched candle holders, due to the structural similarity. Growth of these dendrite branches depends on the DMA-1 dendrite receptor in PVD neurons (Liu and Shen, 2012). What timing mechanisms initiate and terminate the arborization of this complex dendritic structure? To answer these questions, we search for molecules in PVD neurons that may provide intrinsic temporal control of dendritic arborization. We identify two timing regulatory circuits that restrain the growth of PVD dendrites within a set time frame. The initial dendritic outgrowth in PVD neurons is actuated by the precisely timed *lin-4-lin-14* regulatory circuit, which positively and directly modulates the DMA-1 dendrite receptor level through a transcriptional means. The dendritic growth is subsequently slowed down by the precisely timed *lin-28-let-7-lin-41* regulatory circuit, which negatively and indirectly modulates the DMA-1 dendrite receptor level through a posttranscriptional means, as dendritic arborization comes to an end.

## Results

### *lin-4* and *let-7* are temporally expressed in PVD neurons during dendrite arborization

To identify timing mechanisms that restrict PVD dendritic arborization within a defined time period, we sought to identify molecules that are expressed in PVD neurons and whose expression coincides with start and end times of PVD dendritic arborization. We discovered *lin-4* and *let-7* microRNAs to be excellent candidates that fit both criteria. To understand the temporal control of *lin-4* and *let-7* gene expression in PVD neurons, we developed a *lin-4* reporter by fusing 1.9-kb upstream region of mature *lin-4* microRNA to the GFP gene and a *let-7* reporter by fusing 2.9-kb upstream region of mature *let-7* microRNA to the GFP gene and confirmed that they both were stably expressed in PVD (Figure 1). The expression levels of these two reporters in the whole animal at different developmental stages correlated strongly with the whole animal stem-loop reverse transcription polymerase chain reaction (RT-qPCR) quantification of *lin-4* and *let-7* microRNAs, indicating that these two reporters are reliable. Using these reporters, we determined the timing of *lin-4* and *let-7* expression in PVD neurons during dendritic arborization (Figures 1J-1M). The *lin-4* reporter was expressed highly in PVD at the late L2-early L3 stage, when the secondary dendrites start elaborating (Figures 1G, 1H, 1J, and 1L). In contrast, the *let-7* reporter was expressed at relatively low levels in PVD at the L3 stage, but was significantly elevated from the L4 stage onward, when growth of the terminal quaternary dendrites comes to an end (Figures 1G, 1I, 1K, and 1M). These results indicate that *lin-4* and *let-7* are expressed in the right place at the right time to initiate and terminate PVD dendritic arborization, respectively.

**Figure 1.**
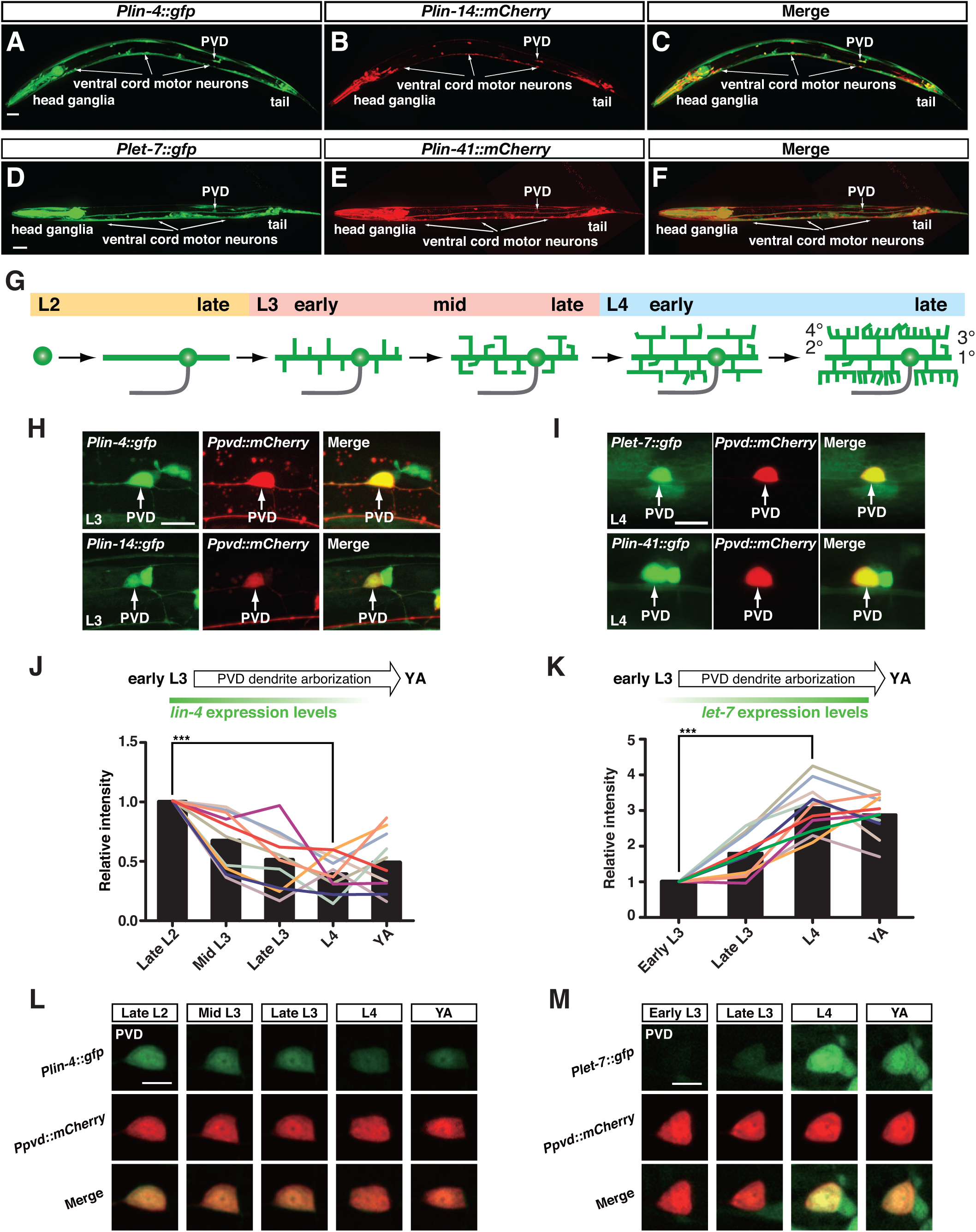
Expression of *lin-4/lin-14* and *let-7/lin-41* regulatory circuits in PVD neurons. (**A-C**) Overlapping expression of *Plin-4::GFP* and *Plin-14::mCherry* reporters in neurons of many regions, such as the head ganglia, ventral nerve cord, tail, and mid-body region, at the third larval (L3) stage. (**D-F**) Overlapping expression of *Plet-7::GFP* and *Plin-41::mCherry* reporters in many neurons in the head ganglia, ventral nerve cord, tail, and mid-body region at the L4 stage. PVD neuron expression was also indicated. Scale bar, 20 µm. (**G**) The timing and steps of PVD dendrite arborization. Schematic drawings of PVD dendrite arbors as they are seen at five different stages of development, late L2 to late L4. All views showing the left-side cells of each PVD pair; anterior is to the left. A single axon (grey color) emerges ventralward from the cell body before traveling anteriorly along the ventral nerve cord. The dendrite processes emerging from the cell body elaborate into highly organized dendrite arbors. (**H**) Both *Plin-4::GFP* and *Plin-14::GFP* reporters are expressed in PVD neurons. (**I**) Detection of both *Plet-7::GFP* and *Plin-41::GFP* expression in PVD neurons. Scale bar, 10 µm. (**J**) Expression of the *Plin-4::GFP* reporter in PVD neurons was assessed at five different stages of development in wild-type animals. (**K**) Expression of the *Plet-7::GFP* reporter in PVD neurons was assessed at four different stages of development in wild-type animals. Each line represents data from a single animal followed over time. Bars represent the average expression intensity of either the *Plin-4::GFP* or the *Plet-7::GFP* reporter measured at each time point. ***p < 0.001 by a Student’s *t*-test. (**L**) Time-lapse imaging of the *Plin-4::GFP* reporter expression in a PVD neuron in wild-type animals at five different stages of development. The *PF49H12.4::mCherry* reporter was used to label PVD neurons. Scale bar, 5 µm. (**M**) Time-lapse imaging of the *Plet-7::GFP* reporter expression in a PVD neuron in wild-type animals at four different stages of development. The *PF49H12.4::mCherry* reporter was used to label PVD neurons. Scale bar, 5 µm.

### The *lin-4*-*lin-14* regulatory circuit initiates dendritic outgrowth in PVD neurons

We previously reported that the *lin-4* microRNA represses the expression of the LIN-14 transcription factor to inhibit AVM axon attraction (Chiu et al., 2014; Zou et al., 2012). Further analysis of *lin-4* and *lin-14* reporters revealed overlapping expression of two genes in many other neurons, including PVD, at the early L3 stage (Figures 1A-1C and 1H), when PVD neurons are sending out the secondary dendrites. We showed that *lin-4(e912)* loss-of-function *(lf)* and *lin-14(n355)* gain-of-function *(gf)* mutants displayed a similar phenotype of limited dendritic outgrowth in PVD neurons (Figures 2A-2C and 2E), suggesting that the *lin-4* microRNA may inhibit the expression of the LIN-14 transcription factor to initiate dendrite outgrowth. Consistent with this interpretation, reduced *lin-14* activity caused opposite effects. In wild-type animals at the mid L3 stage, PVD dendrites can only grow up to the tertiary branch (Figure 2F). However, in *lin-14(n179)* reduction-of-function (*rf*) mutants at the same developmental stage, the mid L3 stage, PVD dendrites can grow up to the quaternary branch and complete the menorah structure (Figure 2F), suggesting precocious dendrite outgrowth. In addition, the number of overlapped tertiary dendrites caused by excessive tertiary dendrite growth was significantly higher in *lin-14(n179rf)* mutants than in wild-type animals at the young adult stage (Figures 2A, 2D, 2G, and S1). Thus, a reduction of function in the *lin-14* gene causes precocious and excessive dendritic outgrowth. Precocious PVD dendritic outgrowth in *lin-14(n179rf)* mutants was not caused by precocious cell fate specification since PVD cells were not specified prematurely in *lin-14(n179rf)* mutants (data not shown). The *lin-14(n179rf)* mutant phenotype of excessive PVD tertiary dendrite growth can be rescued by re-expressing the *lin-14* gene in the PVD neuron, suggesting that *lin-14* acts cell-autonomously in PVD (Figure 2G). To further strengthen the statement that the *lin-4* microRNA targets the *lin-14* transcription factor to initiate dendritic outgrowth in PVD neurons, we tested whether the *lin-14(n179rf)* mutation suppresses the *lin-4(e912lf)* mutant phenotype of limited dendritic outgrowth. We found that this is indeed the case (Figure 2F). *lin-4(lf); lin-14(rf)* double mutants displayed the same precocious dendritic outgrowth phenotype of the *lin-14(n179rf)* single mutant rather than limited dendritic outgrowth phenotypes of the *lin-4(lf)* single mutant (Figure 2F). Taken together, our results indicate that the *lin-4*-*lin-14* regulatory circuit initiates dendritic outgrowth in PVD neurons.

**Figure 2.**
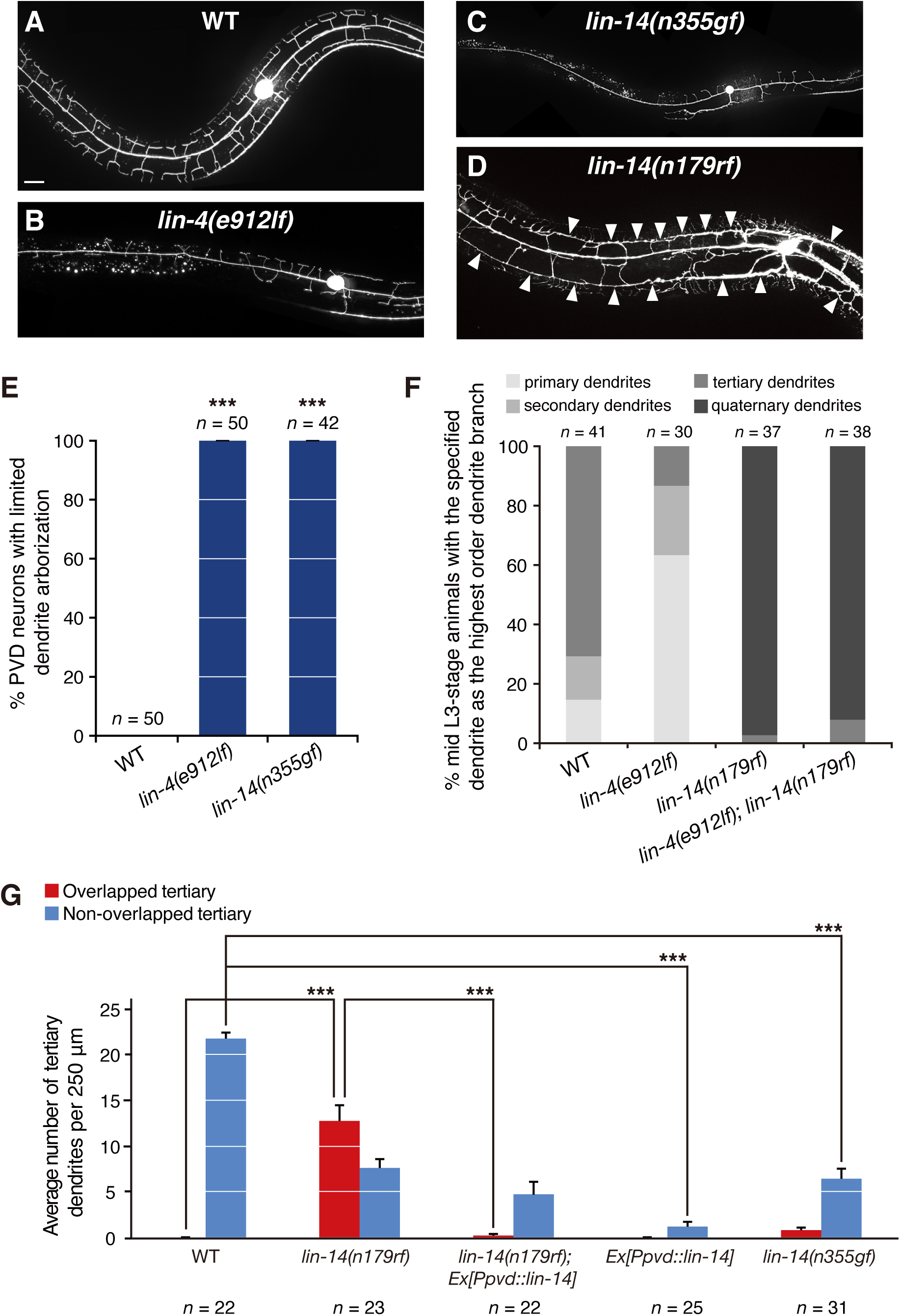
Initiation of dendritic arborization is affected by mutations in the *lin-4-lin-14* regulatory circuit. (**A-D**) Representative images showing extent of dendrite arborization in wild type, *lin-4(e912lf)*, *lin-14(n355gf)*, and *lin-14(n179rf)* mutants. Arrowheads point to contacts between neighboring tertiary dendrites. Scale bar, 20 µm. (**E**) Percentages of PVD neurons at the young adult stage with limited dendrite outgrowth in wild type, *lin-4(e912lf)*, and *lin-14(n355gf)* mutants. Error bars, SEP. ***p < 0.001, relative to wild type, by a two-proportion *Z*-test. (**F**) Percentages of PVD neurons at the mid L3 stage based on the highest order dendrite branch observed in wild type, *lin-4(e912lf)*, *lin-14(n179rf)*, and *lin-4(e912lf)*; *lin-14(n179rf)* mutants. (**G**) Quantification of the number of tertiary branches per 250 µm in the anterior direction from the PVD cell body at the young adult stage in wild type, *lin-14(n179rf)* mutants, *lin-14(n179rf)* mutants carrying the *Pser-2::lin-14* transgene, animals carrying the *Pser-2::lin-14* transgene, and *lin-14(n355gf)* mutants. Tertiary dendrites were divided into two groups: non-overlapped tertiary is defined as those with normal self-avoidance and overlapped tertiary as those with self-avoidance defects. As *lin-14(n179)* is a temperature sensitive allele, *lin-14(n179)* mutants and other strains in this experiment were cultured at a non-permissive temperature, 25°C throughout this study. Error bars, SEM. ***p < 0.001 by a Student’s *t*-test. See also Figure S1.

### The *let-7-lin-41* regulatory circuit slows down dendrite growth in the final stage of dendrite arborization

We previously reported that the *let-7* microRNA represses the expression of the LIN-41 tripartite motif protein to inhibit AVM axon regeneration in older neurons (Chiu and Chang, 2013; Zou et al., 2013). Here, we further studied the expression of *let-7* and *lin-41* reporters and found overlapping expression of two genes in many other neurons, including PVD, at the L4 stage (Figures 1D-1F and 1I), when growth of the terminal quaternary dendrites in PVD comes to an end. We performed laser dendritomy on the primary dendrite in PVD neurons at different developmental stages and found that PVD dendritic growth ability is significantly lower at the adult stage than at the L3 stage (Figures 3A, 3B, and 3E), suggesting that PVD neurons undergo a developmental decline in dendritic growth ability. Our expression analysis showed that the *let-7* expression in PVD was at relatively low levels at the L3 stage, but was significantly elevated from the L4 stage onward, which implicates its contribution to the developmental decline in PVD dendritic growth ability (Figures 1K and 1M). Indeed, the dendritic growth ability in adult *let-7* mutants was indistinguishable from that seen in wild-type animals at an earlier developmental stage, the L3 stage (Figures 3A-3E), suggesting that *let-7* mutations may retard a normal developmental decline in dendritic growth ability. In the young adult stage, while the *let-7(n2853rf)* mutation significantly enhanced, the *lin-41(n2914lf)* mutation significantly reduced dendritic growth ability in PVD neurons (Figure 3F). The *lin-41(n2914lf)* mutant phenotype in PVD dendrites can be rescued by re-expressing the *lin-41* gene in the PVD neuron, suggesting that *lin-41* acts cell-autonomously in PVD. In addition, *lin-41* mutations suppressed the *let-7(n2853rf)* mutant phenotype of enhancing dendrite growth ability, suggesting that the *let-7* microRNA targets the LIN-41 tripartite motif protein to inhibit dendrite growth in PVD neurons (Figure 3F). Together, these results indicate that the *let-7-lin-41* regulatory circuit slows down dendrite growth in the final stage of dendrite arborization.

**Figure 3.**
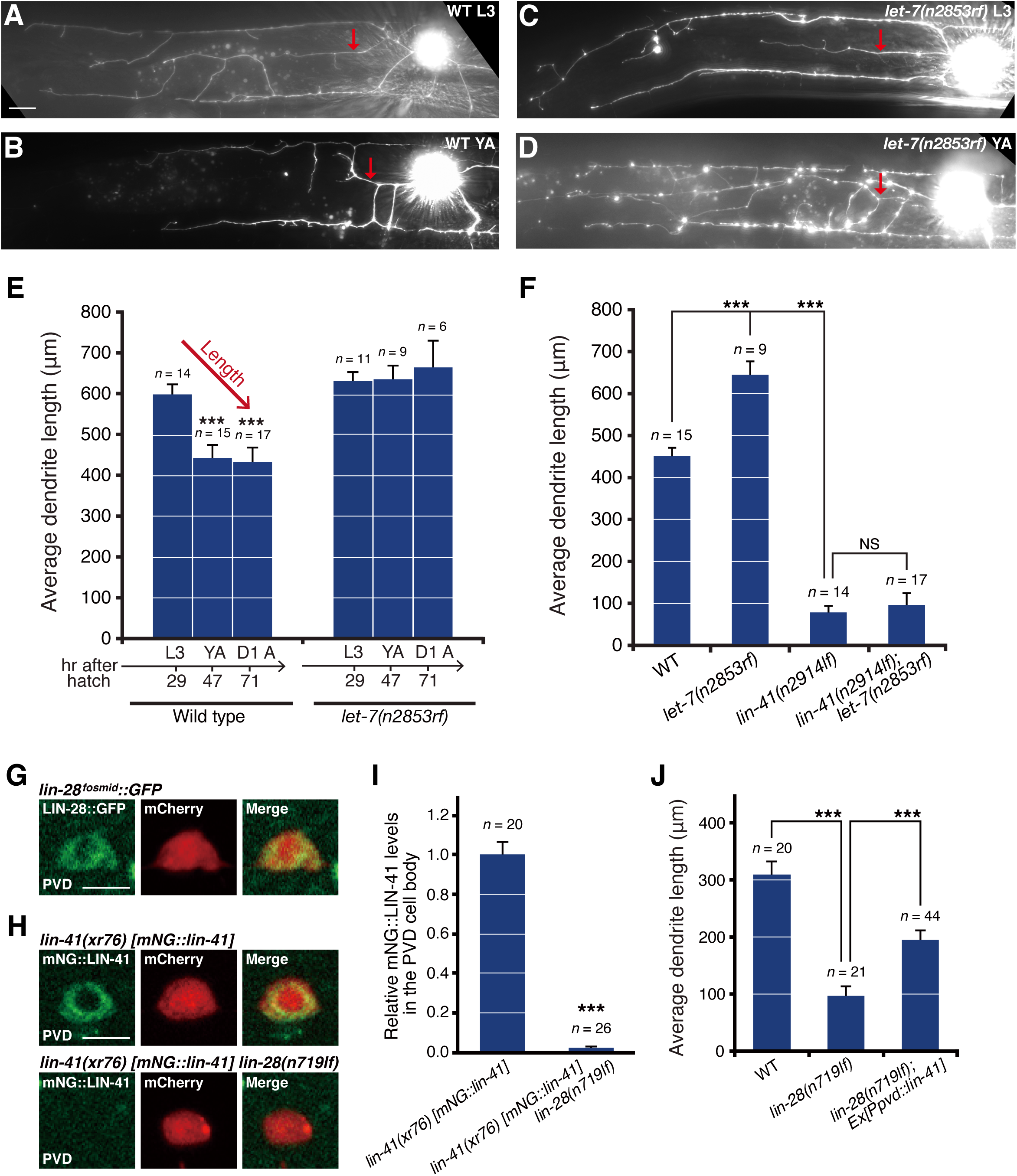
Developmental decline in dendrite growth ability is affected by mutations in the *lin-28-let-7-lin-41* regulatory circuit. (**A-D**) Representative images showing extent of dendrite regrowth 24 hours after dendritomy of the primary dendrite at either the L3 or the young adult stage in wild type (**A, B**) and *let-7(n2853rf)* mutants (**C, D**). PVD dendrites were visualized using the *xrIs37[PF49H12.4::GFP]* marker. Dorsal is up; anterior is to the left. Red arrows indicate lesion sites. Scale bar, 20 µm. (**E**) Average PVD dendrite length regrown in wild type and *let-7(n2853rf)* mutants 24 hours following dendritomy at different stages. Asterisks indicate cases in which later stage animals differ from L3-stage animals at ***p < 0.001 by a Student’s *t*-test. D1 A indicates one day into the adult stage. Error bars indicate SEM. (**F**) Average dendrite length regrown in wild type, *let-7(n2853rf)*, *lin-41(n2914lf)*, and *lin-41(n2914lf); let-7(n2853rf)* mutants. PVD primary dendrites were severed by laser surgery at the young adult stage. Dendrite lengths were measured 24 hours after dendritomy as the actual contour length between the injury site and dendrite termini by tracing dendrites through a 3-dimensional image stack. Asterisks indicate cases in which *let-7* or *lin-41* mutants differ from wild type at ***p < 0.001 by a Student’s *t*-test. NS, not significant. (**G**) Representative images of the expression of LIN-28 proteins in PVD neurons in wild type at the early L3 stage. A *lin-28::GFP* fosmid-based reporter is expressed in PVD neurons. The *Pser-2::mCherry* reporter was used to label PVD neurons. Scale bar, 5 µm. (**H**) Representative images of the expression of endogenous LIN-41 proteins in PVD neurons in wild type and *lin-28(n719lf)* mutants at the early L3 stage. The *Pser-2::mCherry* reporter was used to label PVD neurons. Scale bar, 5 µm. (**I**) Quantification of endogenous LIN-41 proteins based on the mNG::LIN-41 fluorescence intensity in the PVD cell body in wild type and *lin-28(n719lf)* mutants. (**J**) Average dendrite length regrown in wild type, *lin-28(n719lf)* mutants, and *lin-28(n719lf)* mutants overexpressing *lin-41* in PVD neurons. Error bars, SEM. ***p < 0.001 by a Student’s *t*-test. See also Figure S2.

### *lin-28* inhibits the *let-7-lin-41* circuit to regulate the timing of dendrite arborization

Previous studies have shown that the LIN-28 RNA-binding protein blocks maturation of the *let-7* microRNA in both invertebrates and vertebrates (Lehrbach et al., 2009; Vadla et al., 2012; Van Wynsberghe et al., 2011; Viswanathan et al., 2008). We found that a *lin-28::GFP* fosmid-based reporter, which contains the *lin-28* upstream cis-regulatory, exonic, intronic, and downstream cis-regulatory sequences, is expressed in PVD neurons (Figure 3G). To determine whether *lin-28* acts upstream of *let-7* to regulate the timing of PVD dendrite arborization, we first compared the endogenous LIN-41 protein level in PVD neurons between wild type and *lin-28(n719lf)* mutants. The mNG reporter gene was knocked in the endogenous *lin-41* locus to generate a *mNG::lin-41* fusion gene using the CRISPR/Cas9 technology (Shih and Chang, 2021). We found that the *lin-28(n719lf)* mutation significantly reduced LIN-41 protein levels in PVD neurons compared to wild type at the early L3 stage (Figures 3H and 3I), suggesting that *lin-28* inhibits the *let-7-lin-41* circuit in PVD neurons. We performed laser dendritomy on PVD primary dendrites in *lin-28(n719lf)* mutants and observed significantly reduced dendrite growth ability in *lin-28 (n719lf)* mutants compared to wild-type animals 24 hours after surgery (Figure 3J), a phenotype that is opposite to the *let-7(n2853rf)* mutant phenotype of enhanced dendrite growth ability (Figure 3F). Furthermore, *lin-41* overexpression in PVD neurons significantly suppressed the *lin-28(n719lf)* mutant phenotype of reduced dendrite growth ability (Figure 3J). Together, these results support that *lin-28* inhibits the *let-7-lin-41* circuit to regulate the timing of PVD dendrite arborization.

Recent studies in sexually dimorphic nervous system differentiation and male tail tip morphogenesis revealed that the *lep-5* lncRNA brings together LIN-28 and the LEP-2 Makorin to promote ubiquitination and degradation of LIN-28 (Kiontke et al., 2019; Lawson et al., 2019). To determine whether *lep-5* plays a role in regulating the *lin-28-let-7-lin-41* regulatory circuit in PVD neurons, we first analyzed the endogenous LIN-41 protein level in PVD neurons in wild-type animals versus *lep-5(ny28lf)* mutants. We found there is no difference of LIN-41 protein levels in PVD neurons between wild type and *lep-5(ny28lf)* mutants (Figures S2A and S2B). In addition, *lep-5(ny28lf)* mutants displayed similar extent of dendrite growth ability to wild-type animals (Figure S2C). Thus, *lep-5* is unlikely to regulate the timing of PVD dendrite arborization by modulating the *lin-28-let-7-lin-41* circuit.

### The *lin-4* to *let-7* promoter replacement delays dendrite arborization

To further support our conclusion that *lin-4* and *let-7* microRNAs set start and end times for PVD dendrite arborization, we manipulated the timing of their expression by swapping their promoters with each other. We found that delayed expression of *lin-4* (*lin-4* microRNA expressed from the *let-7* promoter from the L3 stage onward) postponed dendrite arborization in PVD neurons (Figure S3). The transgene that expressed the *lin-4* microRNA from the *let-7* promoter (*Plet-7::lin-4*) led to retarded growth in quaternary dendrites (the final order of PVD dendrites) in *lin-4* loss-of-function mutants at early L4 stage (Figures S3A and S3B). However, the growth of quaternary dendrites in these transgenic animals (*lin-4(e912lf); Ex[Plet-7::lin-4]*) was able to catch up later in the adult stage (Figure S3C). To further strengthen this conclusion, we utilized the CRISPR/Cas9 technology to replace the endogenous *lin-4* promoter at the *lin-4* locus with a late-onset *let-7* promoter (Figure 4A’). The repair templates that have been developed recently were used to facilitate the identification of the CRISPR recombinants (Dickinson et al., 2015; Dickinson et al., 2013) *lin-4(xr70)* and *lin-4(xr71)* in which the endogenous *lin-4* promoter has been replaced by a *let-7* promoter. We used stem-loop RT-PCR to globally survey temporal expression of mature *lin-4* microRNA during animal development. Expression of the *lin-4* microRNA in the *lin-4* to *let-7* promoter replacement CRISPR allele was indeed delayed compared to its expression in wild-type animals (Figure 4A). In these CRISPR lines, *lin-4(xr70)* and *lin-4(xr71)*, we observed retarded growth of the quaternary dendrites at the early L4 stage (green dots in Figure 4D compared to 4B of WT; Figure 4F), which was able to catch up later in the adult stage (green dots in Figure 4E compared to 4C of WT; Figures 4G-4I). These results demonstrate that, by manipulating the timing of *lin-4* expression through the *lin-4* to *let-7* promoter replacement, we can delay dendrite arborization.

**Figure 4.**
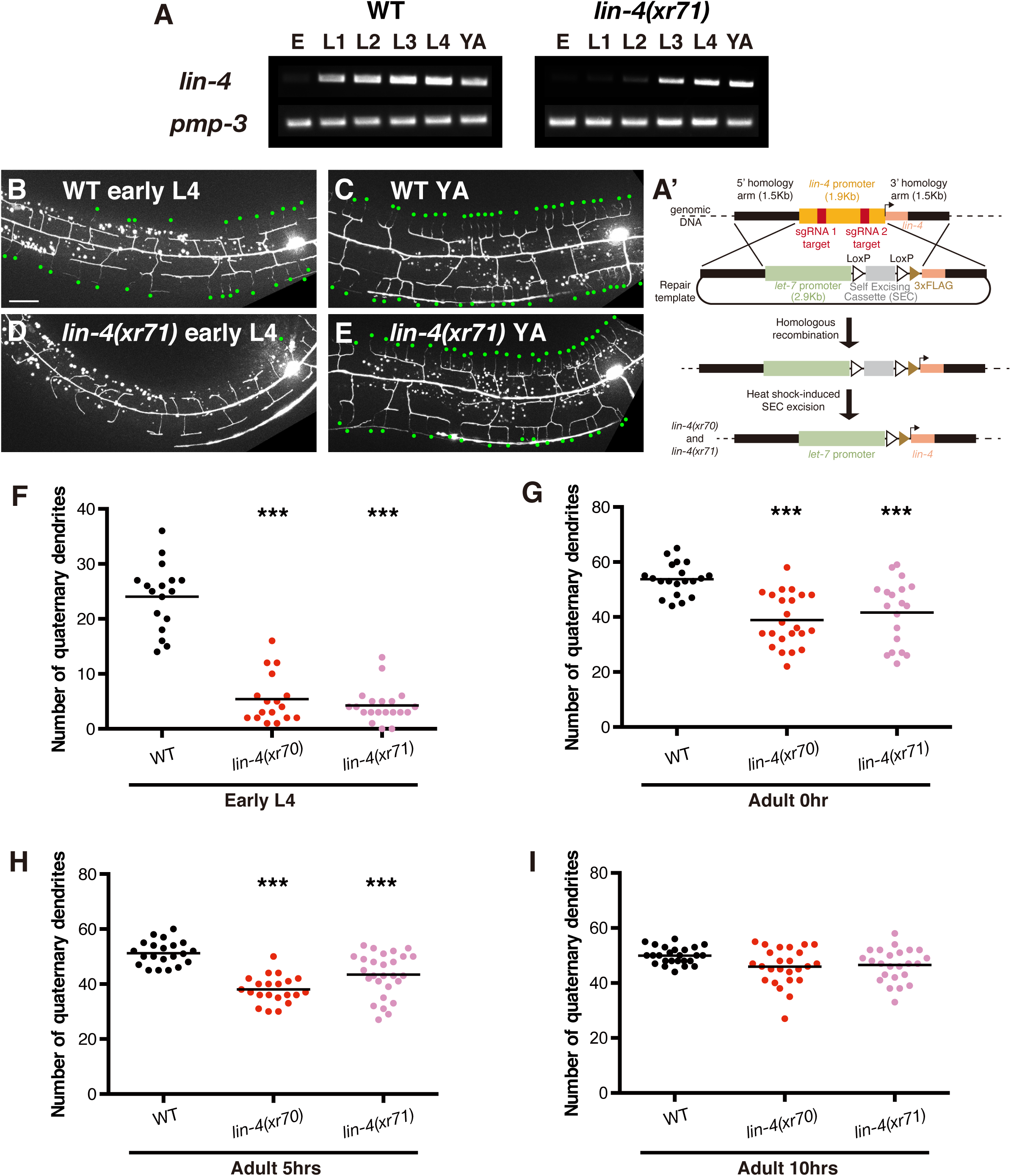
Delayed dendrite arborization by the *lin-4* to *let-7* promoter replacement. Expression of the *lin-4* microRNA by a late-onset *let-7* promoter postponed growth of the quaternary dendrites. (**A**) Stem-loop RT-PCR analysis of RNA isolated from populations of staged animals revealed late onset expression of the *lin-4* microRNA in the *lin-4(xr71)* CRISPR line contrast to early onset expression in wild-type animals. (**A’**) Strategies of promoter replacements by the CRISPR/Cas9 technology. (**B-E**) Representative images showing extent of quaternary dendrite arborization in wild type (**B, C**) and the *lin-4(xr71)* CRISPR line (**D, E**), in which the endogenous *lin-4* promoter has been replaced by the *let-7* promoter. Images were taken at the early L4 (**B, D**) and the young adult (**C, E**) stages. Dorsal is up; anterior is to the left. Green dots indicate quaternary dendrites. Scale bar, 20 µm. (**F-I**) Quantification of the number of quaternary branches per 250 µm in the anterior direction from the PVD cell body at the early L4 stage (**F**) or various time points at the young adult stage (**G-I**) in wild type, *lin-4(xr70)* CRISPR line, and *lin-4(xr71)* CRISPR line. Each dot represents data from a single animal. ***p < 0.001 by a Student’s *t*-test. See also Figure S3.

### The *let-7* to *lin-4* promoter replacement precociously inhibits dendrite growth ability

Conversely, premature expression of *let-7* (*let-7* microRNA expressed from the *lin-4* promoter from the L1 stage onward) precociously inhibited dendritic growth ability in PVD neurons (Figures 5A and 5B). We utilized the CRISPR/Cas9 technology to generate the *let-7(xr67)* CRISPR line in which the endogenous *let-7* promoter at the *let-7* locus has been replaced with an early-onset *lin-4* promoter (Figure 5A’). Stem-loop RT-PCR analysis showed that expression of mature *let-7* microRNA in the *let-7* to *lin-4* promoter replacement CRISPR line was indeed precocious compared to its expression in wild-type animals (Figure 5A). In the *let-7(xr67)* CRISPR line, PVD dendrite growth ability at the L3 stage was significantly lower than that in wild-type animals at the same stage (L3) and similar to that in wild-type animals at an older stage (young adult) (Figure 5B). PVD dendrite growth ability in the *let-7(xr67)* line was further reduced at the young adult stage (Figure 5B). Interestingly, PVD dendrites, both proximal and distal segments to the injured site, in *let-7(xr67)* animals degenerated 24 hours following laser dendritomy at day one into the adult stage (the D1 A stage). Because of this, we were unable to determine the dendrite growth ability at the D1 A stage (Figure 5B). Thus, by manipulating the timing of *let-7* expression through the *let-7* to *lin-4* promoter replacement, we can precociously inhibit dendrite growth ability. Taken together, these findings support a model in which the *lin-4-lin-14* regulatory circuit sets the start time whereas the *let-7-lin-41* regulatory circuit sets the end time for PVD dendrite arborization (Figure 7H).

**Figure 5.**
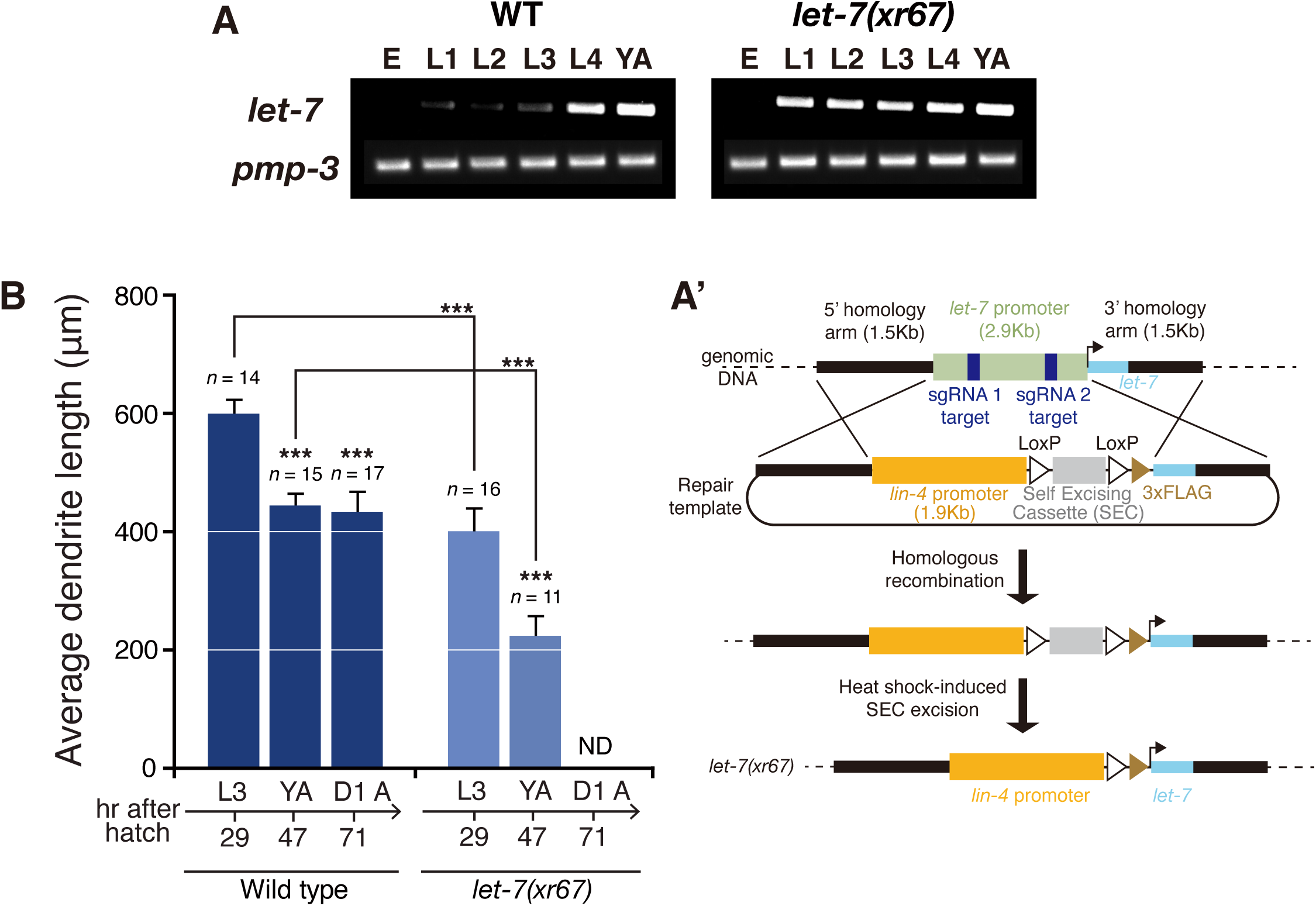
Precocious decline in dendrite growth ability by the *let-7* to *lin-4* promoter replacement. Expression of the *let-7* microRNA by an early-onset *lin-4* promoter precociously reduced dendrite growth ability in PVD neurons. (**A**) Stem-loop RT-PCR analysis of RNA isolated from populations of staged animals revealed early onset expression of the *let-7* microRNA in the *let-7(xr67)* CRISPR line contrast to late onset expression in wild-type animals. (**A’**) Strategies of promoter replacements by the CRISPR/Cas9 technology. (**B**) Average PVD dendrite length regrown in wild type and the *let-7(xr67)* CRISPR line 24 hours following dendritomy of the primary dendrite at different stages. Asterisks indicate cases in which later stage animals differ from L3-stage animals or a comparison between wild type and *let-7(xr67)* is significantly different at ***p < 0.001 by a Student’s *t*-test. D1 A indicates one day into the adult stage. ND represents “not determined”. Error bars indicate SEM.

### *lin-14* and *lin-41* antagonistically regulate the DMA-1 receptor level on PVD dendrites

Two transmembrane ligands, SAX-7 and MNR-1, work together with LECT-2, a secreted ligand, to instruct dendrite arborization in PVD neurons through interactions with the dendrite receptor DMA-1 (Díaz-Balzac et al., 2016; Dong et al., 2013; Liu and Shen, 2012; Salzberg et al., 2013; Zou et al., 2016). SAX-7, MNR-1, LECT-2 and DMA-1 form a multi-protein receptor-ligand signaling complex that directs the growth of stereotyped dendritic branches. Of these four components, only DMA-1 functions cell-autonomously in PVD neurons. One way to control dendrite growth ability during the course of PVD dendrite arborization is through regulation of responsiveness of dendrites to growth signals, which can be accomplished by adjusting the abundance of the receptor on dendrites (Dong et al., 2016). In addition to DMA-1, HPO-30 can function cell-autonomously in PVD neurons as a co-receptor to regulate dendrite arborization by forming a signaling complex with DMA-1 (Smith et al., 2013; Tang et al., 2019; Zou et al., 2018). We ruled out *hpo-30* as a candidate target gene of the two microRNA regulatory circuits since expression of HPO-30 proteins on dendrites was not affected by *lin-14* and *lin-41* mutations, suggesting *hpo-30* is not regulated by *lin-14* and *lin-41* (Figures S4A and S4B). In contrast, expression of DMA-1 proteins on dendrites appeared to be regulated by *lin-14* and *lin-41*. To analyze the endogenous level of DMA proteins on PVD dendrites, we knocked in a GFP reporter after the transmembrane domain of the endogenous *dma-1* gene using the CRISPR/Cas9 technology. This knock-in strain does not show any noticeable defects in PVD dendrites, suggesting that the *gfp* knock-in retains *dma-1* gene function. The endogenous DMA-1 protein level on PVD tertiary dendrites increased in the early stage of dendrite arborization, from the early L3 to early L4 stage, and decreased subsequently in the final stage of dendrite arborization, from the early L4 to young adult stage, which correlates with dynamic change in dendrite growth ability during dendrite arborization (Figure 6A). This observation, combined with our results that two microRNA regulatory circuits set start and end times for PVD dendrite arborization, led us to hypothesize that the *lin-4-lin-14* and the *lin-28-let-7-lin-41* regulatory circuits control the timing of PVD dendrite arborization through antagonistic regulation of the DMA-1 protein level on dendrites (Figure 6B).

**Figure 6.**
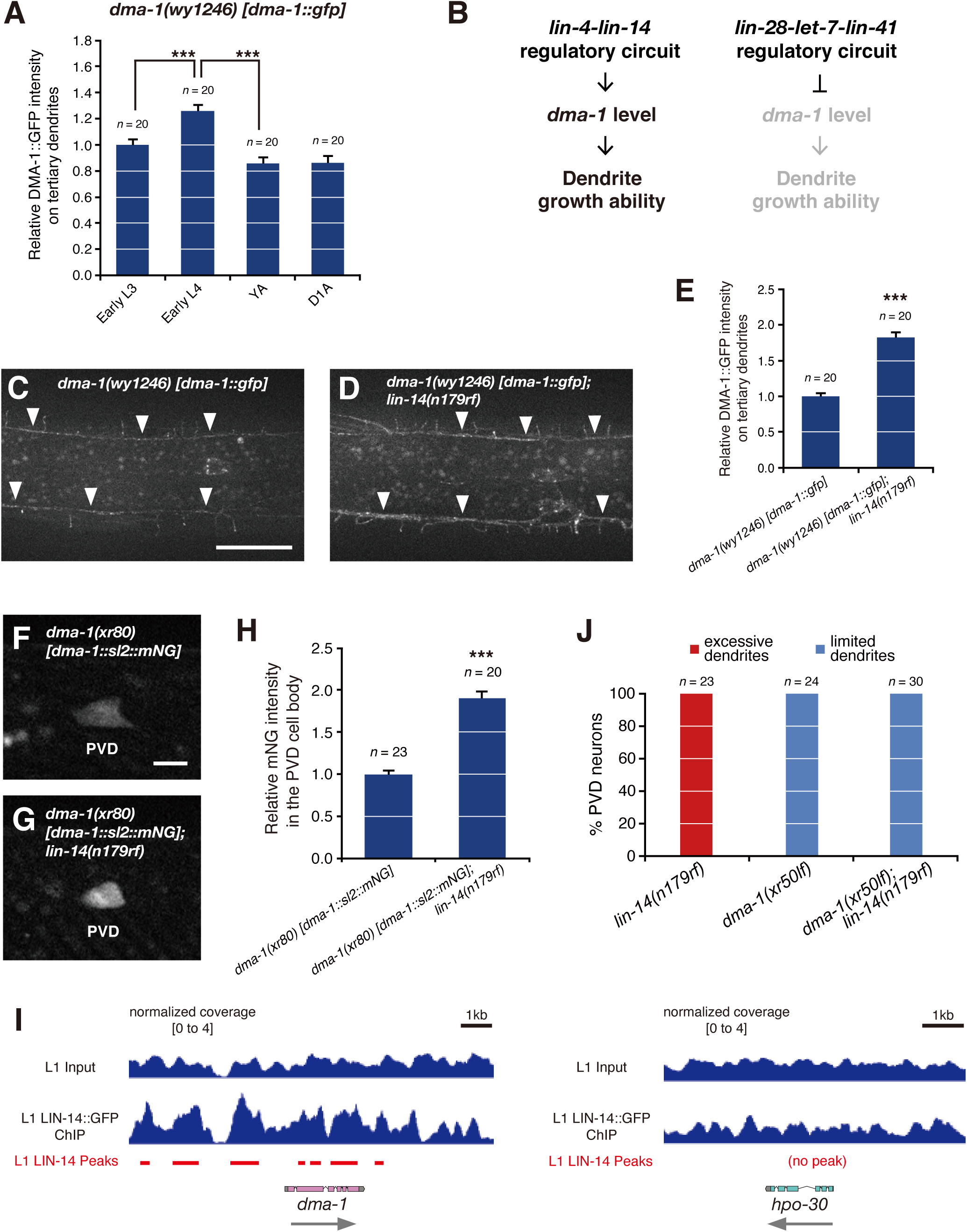
*lin-14* negatively regulates DMA-1 protein levels on PVD dendrites through a transcriptional means. **(A)** Average fluorescence intensity of DMA-1::GFP fusion proteins on PVD tertiary dendrites in wild type (*dma-1(wy1246)*) at four developmental stages. Error bars, SEM. ***p < 0.001 by a Student’s *t*-test. (**B**) Model of temporal control of PVD dendrite growth ability through antagonistic regulation of *dma-1* by two precisely timed microRNA regulatory circuits. (**C, D**) Representative images showing expression of DMA-1::GFP fusion proteins on PVD dendrites in wild type (**C**) and the *lin-14(n179rf)* mutant (**D**). Arrowheads point to tertiary dendrites. Dorsal is up; anterior is to the left. Scale bar, 20 µm. (**E**) Average fluorescence intensity of DMA-1::GFP fusion proteins on PVD tertiary dendrites in wild type (*dma-1(wy1246)*) versus *lin-14(n179rf)* mutants (*dma-1(wy1246); lin-14(n179rf)*). Error bars, SEM. ***p < 0.001 by a Student’s *t*-test. (**F, G**) Representative images showing expression of the SL2-based *dma-1* transcriptional reporter in PVD neurons in wild type (**F**) and the *lin-14(n179rf)* mutant (**G**). Scale bar, 5 µm. (**H**) Average fluorescence intensity of the SL2-based *dma-1* transcriptional reporter in the PVD cell body in wild type (*dma-1(xr80)*) versus *lin-14(n179rf)* mutants (*dma-1(xr80); lin-14(n179rf)*). Error bars, SEM. ***p < 0.001 by a Student’s *t*-test. (**I**) LIN-14 ChIP-seq binding at the *dma-1* gene (left) and the *hpo-30* gene (right). Significant LIN-14 peaks shown in red lines below the ChIP-seq tracks were called by MACS2 (Feng et al., 2012). (**J**) Percentages of PVD neurons at the young adult stage with excessive or limited tertiary dendrite outgrowth in *lin-14(n179rf), dma-1(xr50lf)*, and *dma-1(xr50lf); lin-14(n179rf)* mutants.

To determine whether the *lin-4-lin-14* regulatory circuit controls the timing of PVD dendrite arborization through the regulation of the DMA-1 receptor level on PVD dendrites, we analyzed the level of DMA-1 proteins on tertiary dendrites in *lin-14(n179rf)* mutants versus wild-type animals. We found that the endogenous level of DMA-1 proteins based on the fluorescence intensity of DMA-1::GFP was significantly increased in *lin-14(n179rf)* mutants (Figures 6C-6E), consistent with their higher dendrite growth ability (Figure 2D and 2G). This result suggests that *lin-14* negatively regulates *dma-1* expression in PVD neurons. To understand the mechanism by which *lin-14* negatively regulates the DMA-1 level on PVD dendrites, we engineered a *dma-1* transcriptional reporter by inserting the SL2 trans-splice site and the mNG reporter gene into the 3’ end of the endogenous *dma-1* gene using CRISPR/Cas9 technology. With insertion of this SL2 trans-splice site between *dma-1* and *mNG*, even though they are co-transcribed, they can be translated separately, thus allowing us to read out transcription of the *dma-1* gene based on intensity of the mNG reporter (Tursun et al., 2009). Analysis of the SL2-based *dma-1* transcriptional reporter in the PVD cell body at the early L3 stage showed that *dma-1* transcription was also significantly increased in the *lin-14(n179rf)* mutants compared to wild-type animals (Figures 6F-6H), suggesting that *lin-14* negatively regulates *dma-1* expression in PVD neurons through a transcriptional means. Since the *lin-14* gene encodes a transcription factor, LIN-14 is likely to directly regulate *dma-1* expression. Consistent with this conclusion, the ChIP-seq analysis showed direct binding of LIN-14 transcription factors to immediate upstream and downstream regions of the *dma-1* gene and its gene body (Figure 6I, left). As expected, binding of LIN-14 was not detected around the *hpo-30* gene (Figure 6I, right). To further strengthen the relationship between the *lin-4-lin-14* regulatory circuit and *dma-1* in the context of dendrite growth ability, we analyzed dendrite outgrowth in *dma-1(xr50lf); lin-14(n179rf)* double mutants versus *lin-14(n179rf)* and *dma-1(xr50lf)* single mutants. The *dma-1(xr50)* allele, containing a c-to-t missense mutation that results in a L/F change at aa137, shows limited PVD dendrite outgrowth similar to the null allele. The *dma-1(xr50lf)* mutation significantly suppressed the excessive tertiary dendrite outgrowth caused by the *lin-14(n179rf)* mutation (Figure 6J). Taken together, these results indicate that the *lin-4-lin-14* regulatory circuit promotes dendrite outgrowth through transcriptional up-regulation of the DMA-1 receptor level on dendrites (Figure 7H).

**Figure 7.**
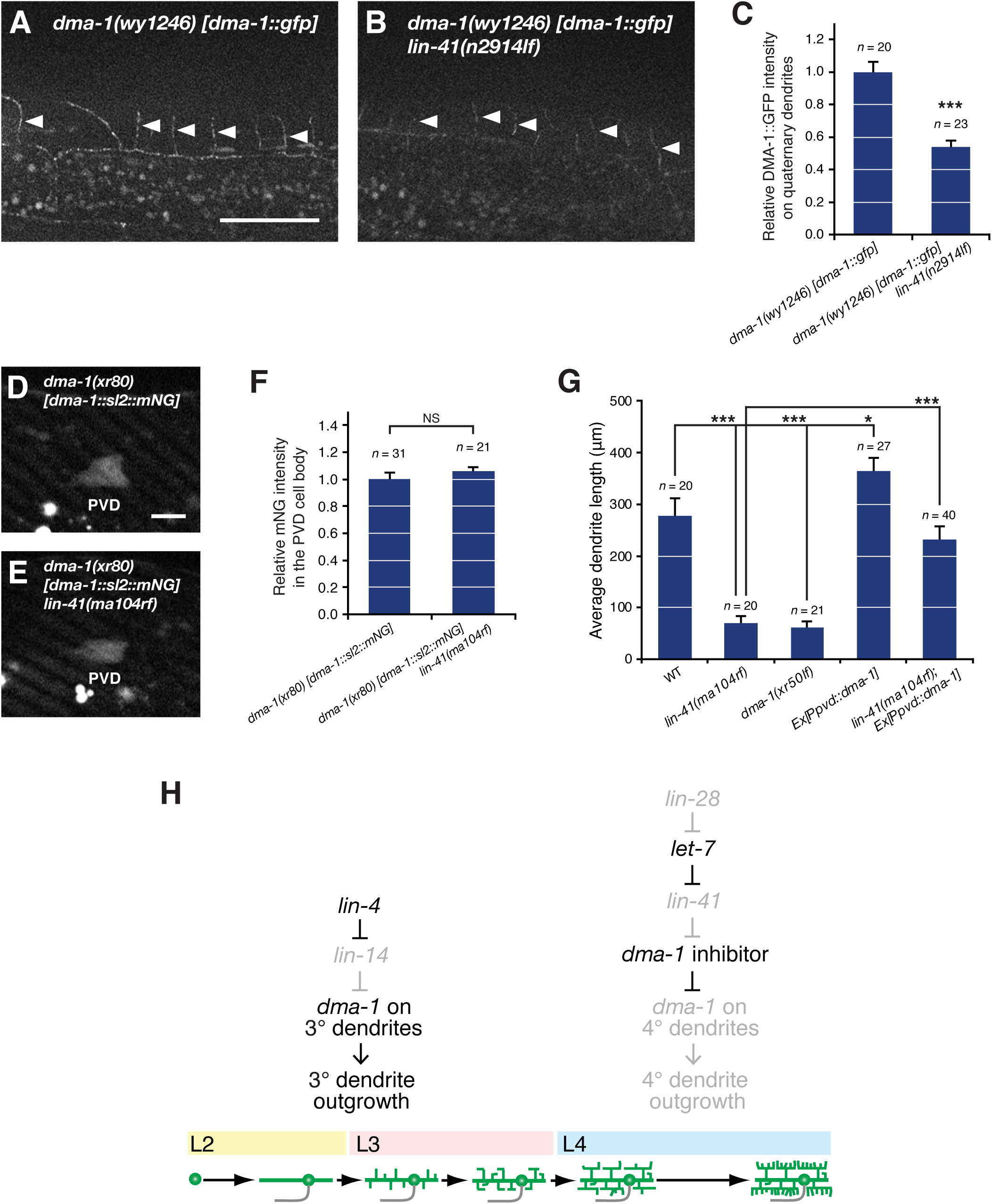
*lin-41* positively regulates DMA-1 protein levels on PVD dendrites through a post-transcriptional means. (**A, B**) Representative images showing expression of DMA-1::GFP fusion proteins on PVD dendrites in wild type (**A**) and the *lin-41(n2914lf)* mutant (**B**). Arrowheads point to quaternary dendrites. Dorsal is up; anterior is to the left. Scale bar, 20 µm. (**C**) Average fluorescence intensity of DMA-1::GFP fusion proteins on PVD quaternary dendrites in wild type (*dma-1(wy1246)*) versus *lin-41(n2914lf)* mutants (*dma-1(wy1246) lin-41(n2914lf)*). Error bars, SEM. ***p < 0.001 by a Student’s *t*-test. (**D, E**) Representative images showing expression of the SL2-based *dma-1* transcriptional reporter in PVD neurons in wild type (**D**) and the *lin-41(ma104rf)* mutant (**E**). Scale bar, 5 µm. (**F**) Average fluorescence intensity of the SL2-based *dma-1* transcriptional reporter in the PVD cell body in wild type (*dma-1(xr80)*) versus *lin-41(ma104rf)* mutants (*dma-1(xr80) lin-41(ma104rf)*). Error bars, SEM. NS, not significant by a Student’s *t*-test. (**G**) Average PVD dendrite length regrown in wild type, *lin-41(ma104rf)* mutants, *dma-1(xr50lf)* mutants, animals overexpressing *dma-1* in PVD neurons, and *lin-41(ma104rf)* mutants overexpressing *dma-1* in PVD neurons 24 hours following dendritomy of the primary dendrite at the young adult stage. Error bars, SEM. *p < 0.05 and ***p < 0.001 by a Student’s *t*-test. (**H**) Model of sculpting dendritic arbors by two precisely timed microRNA regulatory circuits. Initially, the *lin-4* microRNA down-regulates the *lin-14* transcription factor to trigger dendrite arborization. Later, the *let-7* microRNA down-regulates the *lin-41* tripartite motif protein to slow down dendrite growth in the final stage of dendrite arborization. The *lin-14* transcription factor directly represses the *dma-1* gene expression through a transcriptional means whereas the *lin-41* tripartite motif protein indirectly, likely through a *dma-1* inhibitor, promotes the *dma-1* gene expression through a post-transcriptional means.

To determine whether the *lin-28-let-7-lin-41* regulatory circuit also controls the timing of PVD dendrite arborization through the regulation of the DMA-1 receptor level on PVD dendrites, we analyzed the level of DMA-1 proteins on quaternary dendrites in *lin-41(n2914lf)* mutants versus wild-type animals. The endogenous level of DMA-1 proteins was significantly reduced in *lin-41(n2914lf)* mutants (Figures 7A-7C), consistent with their lower dendrite growth ability (Figure 3F). This result suggests that *lin-41* positively regulates the DMA-1 protein level on PVD dendrites. To understand the mechanism by which *lin-41* positively regulates *dma-1* expression, we analyzed expression of the SL2- based *dma-1* transcriptional reporter in the PVD cell body in *lin-41(ma104rf)* mutants versus wild type at the early L3 stage. The *lin-41(ma104rf)* mutation did not affect *dma-1* transcription, suggesting that *lin-41* positively regulates the DMA-1 protein level on PVD dendrites through a post-transcriptional means (Figures 7D-7F). Considering that the known role of LIN-41, either as an RNA-binding protein or a ubiquitin ligase, is to negatively regulate target gene expressions (Aeschimann et al., 2017; Pereira et al., 2019; Slack et al., 2000; Zou et al., 2013), LIN-41 most likely positively regulates the DMA-1 level through an indirect mechanism such as negative regulation of a *dma-1* inhibitor (Figure 7H). To further strengthen the relationship between the *lin-28-let-7-lin-41* regulatory circuit and *dma-1* in the context of dendrite growth ability, we performed laser dendritomy on PVD neurons in *lin-41(ma104rf)* mutants with or without *dma-1* overexpression in PVD neurons. *dma-1* overexpression in PVD neurons displayed enhanced dendrite growth ability, which is opposite to reduced dendrite growth ability caused by the *dma-1(xr50lf)* mutation (Figure 7G). *dma-1* overexpression in PVD neurons significantly suppressed the reduced dendrite growth ability caused by the *lin-41(ma104rf)* mutation (Figure 7G). Thus, this result indicates that the *lin-28-let-7*-*lin-41* regulatory circuit inhibits dendrite growth ability through post-transcriptional down-regulation of the DMA-1 receptor level on dendrites (Figure 7H). Taken together, our findings support a model in which the *lin-4-lin-14* and the *lin-28-let-7-lin-41* regulatory circuits control the timing of PVD dendrite arborization through antagonistic regulation of the DMA-1 receptor level on PVD dendrites (Figure 7H). The LIN-14 transcription factor acts directly to transcriptionally repress *dma-1* expression whereas the LIN-41 tripartite motif protein acts indirectly to post-transcriptionally promote *dma-1* expression in PVD neurons.

A candidate gene that can act between *lin-41* and *dma-1* in PVD neurons is the *lin-29* zinc finger transcription factor as it is a known *lin-41* target gene and is expressed in PVD nuclei (Figure S5A). *lin-29* has been reported to be negatively regulated by *lin-41* in certain cellular contexts in *C. elegans* (Aeschimann et al., 2017; Pereira et al., 2019; Slack et al., 2000; Zou et al., 2013). Although the LIN-29 transcription factor is expressed in PVD nuclei strongly from the L4 stage onward (Figures S5B), the *lin-29(n333lf)* mutation did not affect *dma-1* transcription (Figure S5C), suggesting that *lin-29* is unlikely to act between *lin-41* and *dma-1* in PVD neurons. To assess whether *lin-29* acts as a *dma-1* inhibitor in PVD neurons (Figure 7H), we examined the effect of *lin-29* deficiency on PVD dendrite growth ability. Due to the fact that *lin-29* is required in not only PVD neurons but also surrounding hypodermal tissues that envelop PVD dendrites, we used the ZIF-1-mediated protein degradation system to deplete LIN-29 proteins specifically in PVD neurons (Armenti et al., 2014; Wang et al., 2017). This PVD-specific LIN-29 knockdown did not affect PVD dendrite growth ability (Figure S5D), further supporting the notion that *lin-29* is unlikely to act between *lin-41* and *dma-1* to control the timing of PVD dendrite arborization.

## Discussion

PVD neurons lose dendrite growth ability as they age, but it is not known why. One theory is that the developmental decline in dendrite growth ability provides a switch in the internal state from long-range exploratory growth to short-range dendritic targeting. In this report, we show that two microRNA regulatory circuits are used in timing dendritic arborization in postmitotic PVD neurons, restricting dendrite growth within a defined time frame. The precisely timed *lin-4-lin-14* regulatory circuit sets off initial dendrite outgrowth until the L4 stage, at which the precisely timed *lin-28-let-7-lin-41* regulatory circuit decelerates dendrite growth as terminal dendrite branches are reaching final targets (Figure 7H). These two regulatory circuits control the timing of PVD dendrite arborization through opposed regulation of the DMA-1 receptor level on PVD dendrites (Figure 7H).

The neuronal timers that restrain dendrite arborization within a specific time window during neural circuit formation are poorly understood and remain a mystery. In this study, we uncover two microRNA regulatory circuits that temporally control the choreography of dendrite arborization in PVD neurons. Our study illustrates how neuronal timers regulate the intrinsic potential of dendrite growth. From this point forward, our goal is to describe the required timing mechanisms in dendritic arborization at a resolution that ultimately will allow us to reconstitute the process. Further dissection of genetic networks that regulate the timing of *lin-4* and *let-7* expression, and identification of the downstream target of the LIN-41 tripartite motif protein that regulates the DMA-1 receptor level on dendrites will be indispensable to achieving this goal.

Here, we show that the *lin-4-lin-14* regulatory circuit initiates PVD dendrite arborization through up-regulation of the DMA-1 receptor level on dendrites. The LIN-14 transcription factor most likely binds to the cis-regulatory region of *dma-1* to negatively regulate *dma-1* transcription (Figures 6F-6I). We also report that the *lin-28-let-7-lin-41* regulatory circuit slows down PVD dendrite arborization as it comes to an end through down-regulation of the DMA-1 receptor level on dendrites. The positive regulation of the DMA-1 level by LIN-41 is likely indirect since LIN-41 is known to negatively regulate its target genes (Aeschimann et al., 2017; Pereira et al., 2019; Slack et al., 2000; Zou et al., 2013). The LIN-41 tripartite motif protein is proposed to negatively regulate a *dma-1* inhibitor to post-transcriptionally promote the DMA-1 receptor level on PVD dendrites (Figures 7D-7F). Although the *lin-29* transcription factor, a known direct target of LIN-41 in certain cellular contexts (Aeschimann et al., 2017; Pereira et al., 2019; Slack et al., 2000; Zou et al., 2013), is expressed in PVD neurons (Figures S5A and S5B), *lin-29* does not regulate *dma-1* transcription and dendrite growth ability (Figures S5C and S5D), suggesting that *lin-29* is unlikely to regulate the timing of PVD dendrite arborization. Thus, further studies are required to understand the mechanism by which LIN-41 regulates the DMA-1 receptor level on PVD dendrites.

In the *let-7(xr67)* CRISPR line, the *let-7* microRNA expressed from the *lin-4* promoter does not block PVD dendrite growth in early development. This could be due to many different reasons. For example, the *Plin-4::let-7* genomic configuration may not be able to support expression and processing of the primary *let-7* microRNA as efficiently as the *Plin-4::lin-4* genomic configuration supports expression and processing of the primary *lin-4* microRNA. Alternatively, between the *lin-4-lin-14* and the *let-7-lin-41* regulatory circuits, the *lin-4-lin-14* regulatory circuit could impact the DMA-1 receptor level to a larger extent than the *let-7-lin-41* regulatory circuit.

It remains to be seen whether the timing of dendrite arborization by *lin-4* and *let-7* microRNA regulatory circuits can be extended beyond PVD neurons, especially knowing that these two timing microRNAs are expressed broadly in many neurons in *C. elegans* (Figures 1A-1F) (Shih and Chang, 2021; Zou et al., 2012; Zou et al., 2013). Since *let-7* and *lin-4* microRNAs are evolutionarily conserved, it is possible that these microRNA regulatory circuits control the timing of dendrite arborization in other organisms as well. This possibility is supported by the findings that *let-7* is expressed in the nervous system in various species, including *C. elegans*, *Drosophila*, mouse, and human (Fairchild et al., 2019; Pasquinelli et al., 2000; Sempere et al., 2002; Zou et al., 2013) and that *miR-125,* the *lin-4* homolog, functions in the *Drosophila* and mouse nervous system (Åkerblom et al., 2014; Wu et al., 2012). Although the timing of dendrite arborization is less understood in vertebrates, untimely dendrite arborization likely would lead to neurodevelopmental and psychiatric disorders that are frequently associated with defective dendritic arbors (Koleske, 2013). Future studies in other systems could reveal whether temporal control of dendrite arborization by the two microRNA regulatory circuits is evolutionarily conserved.

In this study, we discover a role for the *lin-4-lin-14* and the *lin-28-let-7-lin-41* regulatory circuits in temporal control of PVD dendrite arborization. We do not rule out the possibility that these microRNA regulatory circuits may also be involved in other differentiation events in PVD neurons. Indeed, using a PVD neuronal marker to assess its differentiation state in *lin-14(n179rf)* mutants versus wild-type animals, we found that PVD neurons in *lin-14(n179rf)* mutants exhibit precocious differentiation (Figure S6), suggesting that timing of many differentiation events in PVD neurons may also be controlled by the *lin-4-lin-14* regulatory circuit.

## Methods

### Strains and plasmids

*C. elegans* strains were cultured using standard methods (Brenner, 1974). All strains were grown at 20°C, except for experiments that involved the *lin-14(n179ts)* allele, which was grown at 25°C. Standard protocol was used for the plasmid constructions. Strains and plasmids used in this study are listed in Tables S1 and S2.

### Transgenic animals

Germline transformation of *C. elegans* was performed using standard techniques (Mello and Fire, 1995). For example, the *Plin-41::GFP* transgene was injected at 50 ng/µl along with the coinjection marker *Podr-1::RFP* at 50 ng/µl. Transgenic lines were maintained by following the *Podr-1::RFP* fluorescence.

### CRISPR/Cas9 genome editing

We generated the *lin-4(xr70)*, *lin-4(xr71)*, *let-7(xr67)*, *let-7(xr68)*, and *dma-1(xr80)* CRISPR lines using the CRISPR/Cas9 genome editing technology. CRISPR recombinants were identified using a self-excising drug selection cassette as previously described (Dickinson et al., 2015; Dickinson et al., 2013). The strategies are illustrated in the diagrams in Figures 4A’ and 5A’.

### Stem-loop reverse transcription-PCR

We quantified the mature microRNA level by modifying the microRNA assay developed previously (Chen et al., 2005). Equal amounts of the RNA preparation from staged wild type, *lin-4(xr71)*, and *let-7(xr67)* animals were used for RT-PCR amplification of mature microRNAs and of *pmp-3* transcripts. Reverse transcription reactions contained purified total RNA, 50 nM stem-loop RT primer, 1X RT first strand buffer, 0.25 mM each of dNTPs, 10 mM MgCl_2_, 0.1 M DTT, 200 U SuperScript III reverse transcriptase, and 40 U RNase inhibitor. The mixture of RNA template, dNTPs, and RT primer was incubated for 5 min at 65°C. The mixture was then placed on ice for at least 1 min before adding RT buffer, DTT, MgCl_2_, SuperScript III, and RNase inhibitor. The reaction was incubated for 50 min at 50°C before heat inactivation at 85°C for 5 min. PCR was conducted using 0.25 µl RT products as template in 20 µl PCR for 17 cycles.

RT_primer_lin-4 TCAACTGGTGTCGTGGAGTCGGCAATTCAGTTGAGTCACACTT
lin-4_F CGGCGGTCCCTGAGACCTCAA
RT_primer_let-7 CTCAACTGGTGTCGTGGAGTCGGCAATTCAGTTGAGAACTATAC
let-7_F CGGCGGTGAGGTAGTAGGTTGT
Universal reverse primer CTGGTGTCGTGGAGTCGGCAATTC

pmp-3-F TGGCCGGATGATGGTGTCGC
pmp-3-R ACGAACAATGCCAAAGGCCAGC

### Laser dendritomy

Animals were mounted on 2% agarose pads and anesthetized with 5 mM sodium azide, the lowest possible concentration to keep adult animals immobilized. Laser dendritomy was performed on PVD primary dendrites using either a cavity-dumped Ti:sapphire laser oscillator (Cascade Laser, KMLabs Inc.) (Chiu et al., 2018; Zou et al., 2013), which generates laser pulses ∼100 fs in duration and 200 kHz in repetition rate, or a MicroPoint Laser Ablation System (Andor/Oxford Instruments), which generates 337 nm laser pulses 2-6 ns in duration. The laser pulses were tightly-focused onto targeted primary dendrites using a Plan Apo VC 100x, 1.4 NA oil-immersion objective on a Nikon ECLIPSE Ti microscope. Successful laser dendritomy was confirmed by visualizing the targeted area immediately after surgery. Worms were recovered within 10 minutes of sodium azide treatment and placed on fresh plates with bacterial food.

### Quantification of dendrite lengths

PVD neurons in recovered worms were imaged 24 hours after dendritomy and the dendrite regrowth was quantified. Dendrite lengths were calculated as the actual contour length between the injury site and dendrite termini measured along the cylindrical surface of each worm by tracing dendrites through a 3-dimensional image stack. P values for the length measurements were calculated using a Student’s *t*-test.

### Fluorescence microscopy

Animals were mounted on 2% agarose pads and anesthetized with 20 mM sodium azide. Fluorescence microscopy was performed using either a Plan-Apochromat 60x, 1.4 NA objective on a Zeiss Axio Imager M2 microscope with a Hamamatsu ORCA-Flash4.0 LT+ camera, or a Plan Apo VC 60x, 1.4 NA objective on a Nikon ECLIPSE Ti microscope with a Hamamatsu ORCA-ER camera. The morphology of neuronal cell bodies and dendrites was based on high-magnification Z-stacks.

### ChIP-seq and analysis

The C-terminally GFP-tagged *lin-14* (*lin14(cc2841)[lin-14::gfp]*) strain was used for the ChIP-seq analysis. Around 600,000 L1 animals were collected and fixed with 2% formaldehyde for 15 min at room temperature. The ChIP-seq analysis was performed as previously described with modifications (Sun and Hobert, 2021). The immunoprecipitated DNA was purified using Ampure XP beads (A63881) according to the manufacturer’s instructions, and used to generate sequencing library using Ovation Ultralow System V2 (Tecan) according to the manufacturer’s instructions. The libraries were sequenced on Illumina NextSeq 500 machines with 75 bp single-end reads. After initial quality check, the reads were mapped to WS220 using BWA (Li and Durbin, 2009) and filtered using SAMtools (Li et al., 2009). Peaks were called using MACS2 (Feng et al., 2012). All peaks and differential binding sites were annotated and assigned to the nearest gene using ChIPseeker (Yu et al., 2015).

### Statistical analysis

Average data of dendrite number, dendrite length, and relative fluorescent reporter intensity are presented as means ± SEM. Data of % PVD neurons with excessive or limited dendrites are presented as proportions ± SEP. Statistical analyses were carried out by Student’s *t*-tests or two-proportion *Z*-tests using GraphPad Prism 7.0 or the Primer of Biostatistics software. The statistical test used for each panel is described in the figure legend. Sample sizes for experiments are shown in the respective panels. In all figures, *n* represents number of animals examined.

## Acknowledgments

This work was funded by grants from the March of Dimes Foundation (C.C.), the Whitehall Foundation Research Award (C.C.), the National Science Foundation (IOS-1455758 to C.C.), the National Institute of General Medical Sciences of the National Institutes of Health (R01GM111320 to C.C.), and the Howard Hughes Medical Institute (K.S.). We thank Hui Chiu for critical reading the manuscript, providing critical initial data, and making figures, Daniel J. Dickinson and Bob Goldstein for providing reagents and CRISPR protocols, David H.A. Fitch for the *lep-5(ny28)* mutant allele, Oliver Hobert for the *lin-29(xe61)* reporter strain and unpublished data, Daniel D. Shaye and Chiou-Fen Chuang for constructs, Evguenia Ivakhnitskaia for technical assistance and critical reading the manuscript, Rui Xiong for technical assistance, the *Caenorhabditis* Genetics Center for strains, and the WormBase for readily accessible information.

## Author Contributions

N.S. conceived, designed, performed, analyzed experiments, made constructs, and drafted the article. Y.Z. conceived, designed, performed, analyzed experiments, and made constructs. H.S. designed, performed, analyzed the ChIP-seq, and contributed unpublished essential data. K.E. made the *dma-1(wy1246)* [*dma-1::gfp*] CRISPR line and contributed unpublished essential data and reagents. M.S. made the promoter replacement CRISPR lines, performed, and analyzed experiments. K.S. conceived experiments, contributed unpublished essential data and reagents, and drafted the article. C.C. conceived, designed, analyzed and interpreted data, and drafted the article.

