## Supplementary Information for "Two intrinsic timing mechanisms set start and end times for dendritic arborization of a nociceptive neuron"

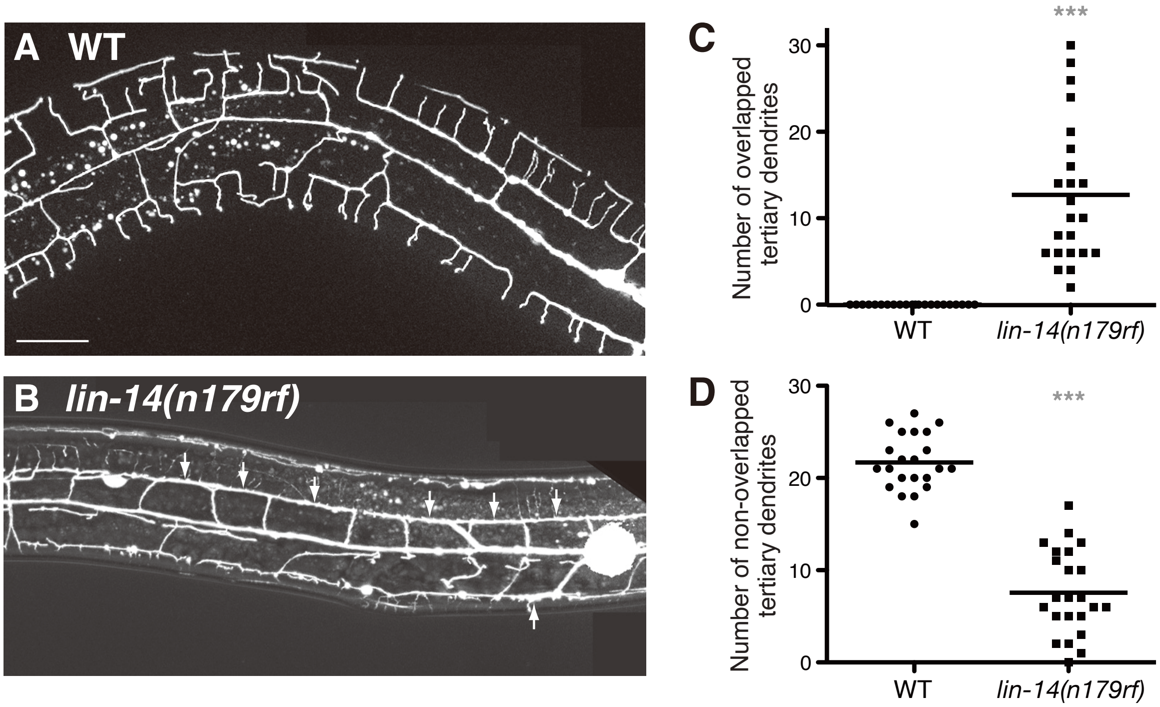


**Figure S1.** ***lin-14(n179rf)*** **mutations caused excessive PVD dendrite outgrowth, related to Figure 2.** (**A, B**) Images of PVD dendrites in wild type (**A**) and *lin-14(n179rf)* mutants (**B**). Dorsal is up; anterior is to the left. Scale bar, 20 μm. *lin-14(n179rf)* mutants displayed excessive tertiary dendrites. Arrows point to contacts between neighboring tertiary dendrites, which are absent in wild-type animals. (**C**) Quantification of the number of overlapped tertiary dendrites in wild type versus *lin-14(n179rf)* mutants. (**D**) Quantification of the number of non-overlapped tertiary dendrites in wild type versus *lin-14(n179rf)* mutants. Each dot represents data from a single animal. ***$p < 0.001$ by a Student's *t*-test.

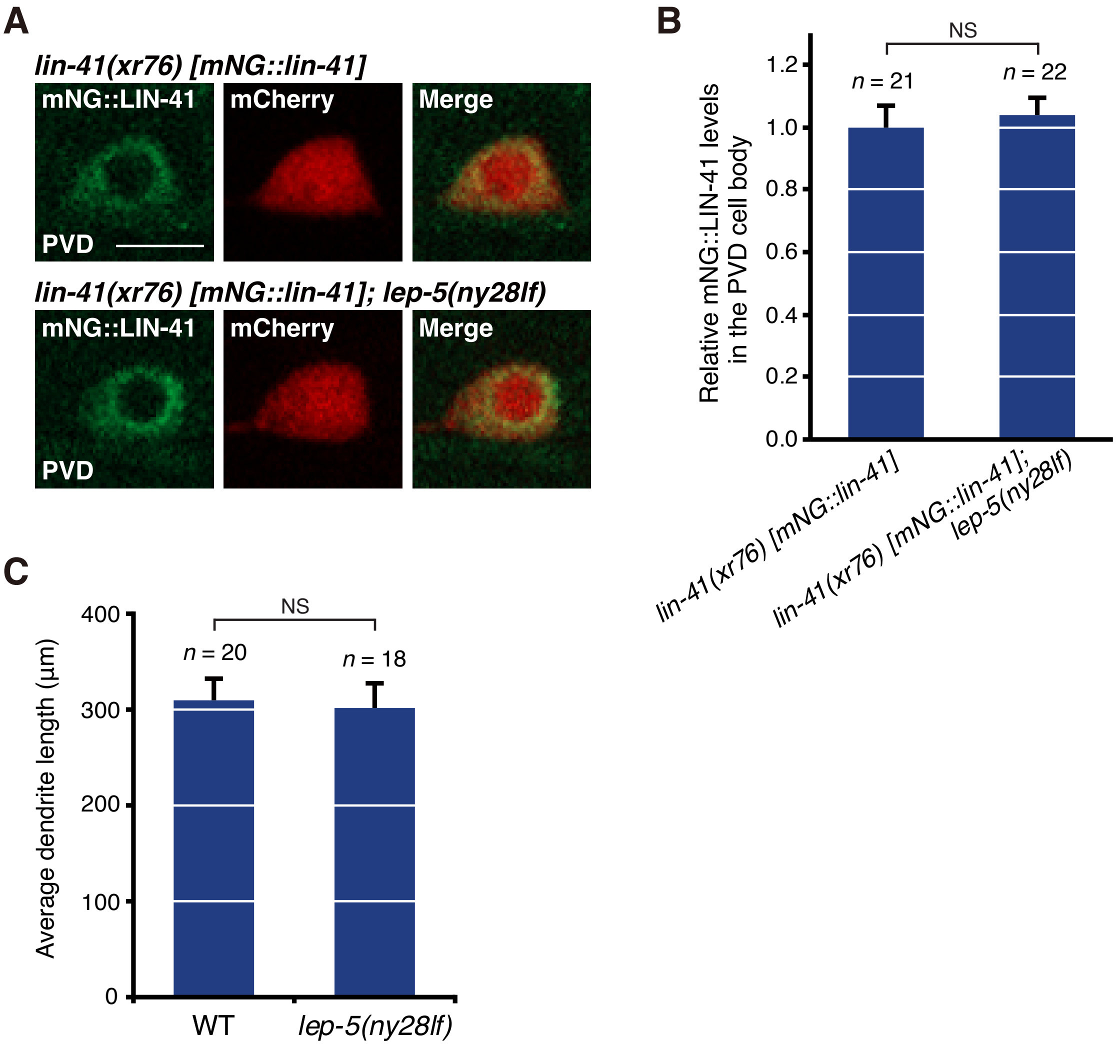


**Figure S2.** ***lep-5*** **does not regulate dendrite growth ability, related to Figure 3.** (**A**) Representative images of the expression of endogenous LIN-41 proteins in PVD neurons in wild type and *lep-5(ny28lf)* mutants at the early L3 stage. The *Pser-2::mCherry* reporter was used to label PVD neurons. Scale bar, 5 μm. (**B**) Quantification of endogenous LIN-41 proteins based on the fluorescence intensity in the PVD cell body in wild type and *lep-5(ny28lf)* mutants. (**C**) Average dendrite length regrown in wild type and *lep-5(ny28lf)* mutants 24 hours following dendritomy of the primary dendrite at the young adult stage. Error bars, SEM. NS, not significant by a Student's *t*-test.

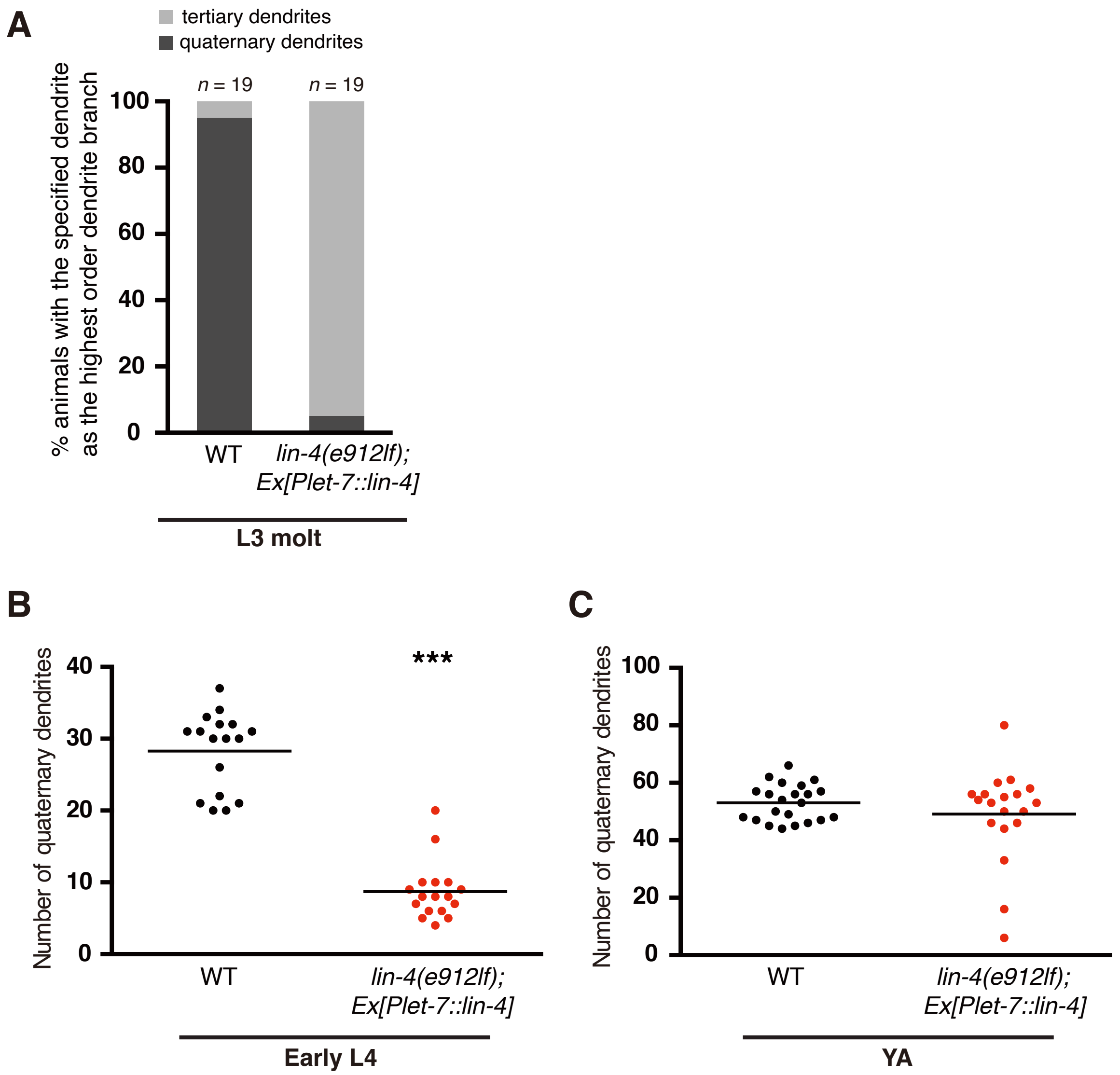


**Figure S3. Delayed dendrite arborization by the *lin-4* to *let-7* promoter replacement, related to Figure 4.** (**A**) Re-expression of the *lin-4* microRNA in *lin-4(e912lf)* mutants using a late-onset *let-7* promoter postponed growth of the quaternary dendrites. (**B, C**) Quantification of the number of quaternary branches per 250 μm in the anterior direction from the PVD cell body at the early L4 stage (**B**) or the young adult stage (**C**) in wild type and *lin-4(e912lf)* mutants carrying the *Plet-7::lin-4* transgene. ***$p < 0.001$ by a Student's *t*-test.

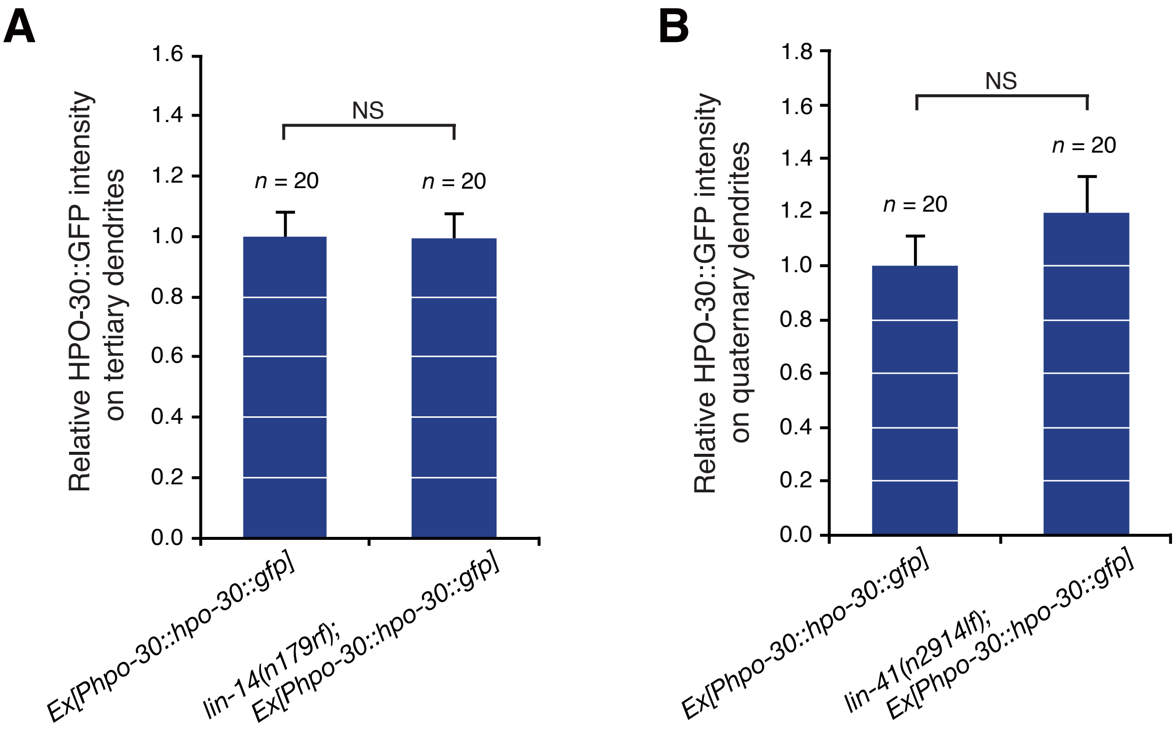


**Figure S4.** ***hpo-30*** **is not regulated by** ***lin-14*** **or** ***lin-41*****.** (**A**) Average fluorescence intensity of the *Phpo-30::hpo-30::gfp* reporter on PVD tertiary dendrites in the wild-type versus *lin-14(n179rf)*-mutant background. (**B**) Average fluorescence intensity of the *Phpo-30::hpo-30::gfp* reporter on PVD quaternary dendrites in the wild-type versus *lin-41(n2914lf)*-mutant background. Error bars, SEM. NS, not significant by a Student's *t*-test.

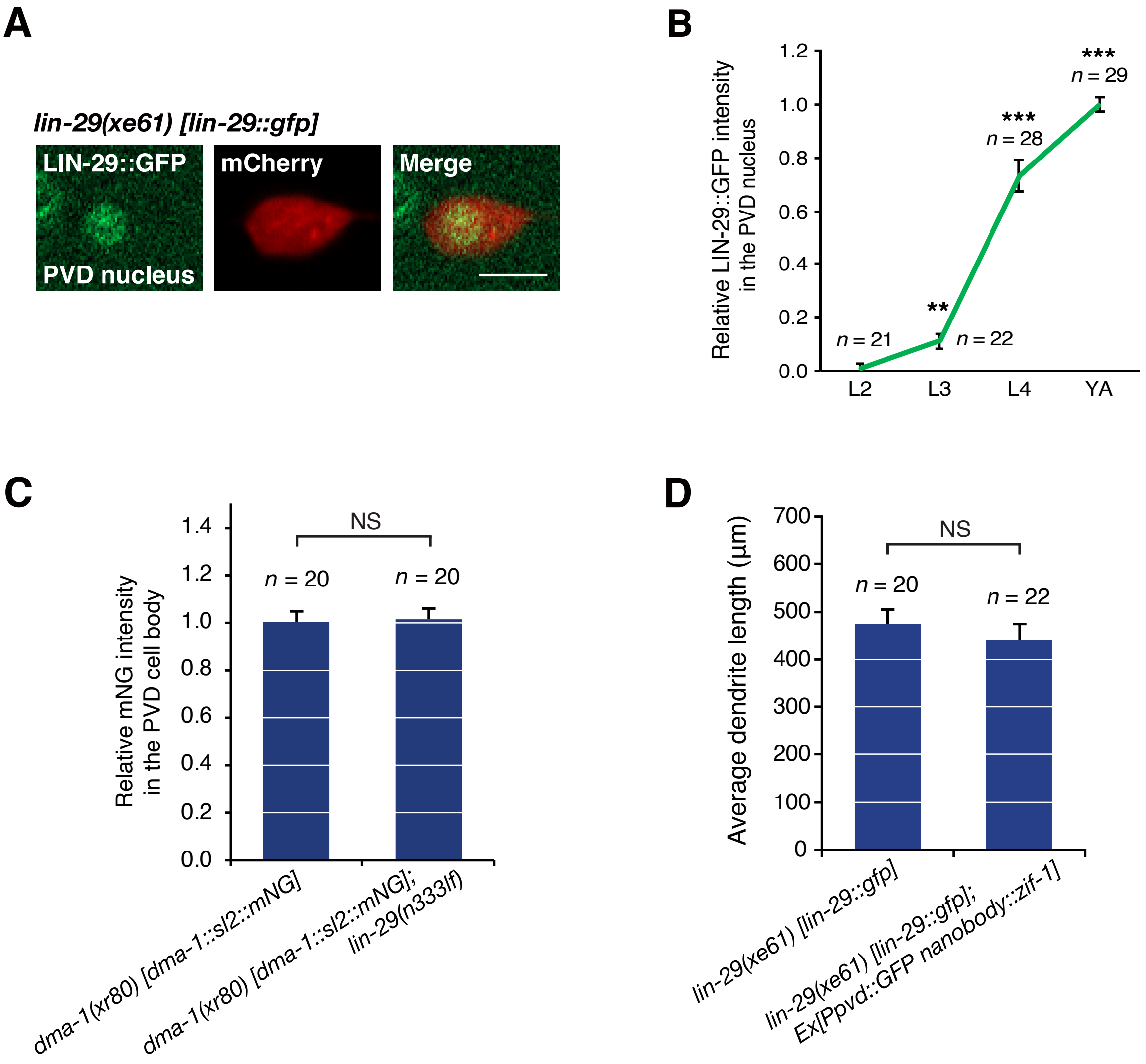


**Figure S5. *lin-29* is not involved in regulating dendrite growth ability despite its expression in PVD neurons.** (**A**) Representative images of the expression of endogenous LIN-29 proteins in the PVD nucleus in wild type at the young adult stage. The *Pser-2::mCherry* reporter was used to label PVD neurons. Scale bar, 5 µm. (**B**) The endogenous LIN-29 protein is temporally regulated in PVD neurons. Average fluorescence intensity of LIN-29::GFP proteins at four different developmental stages. Error bars, SEM. **p < 0.01 and ***p < 0.001, relative to the preceding stage, by a Student's *t*-test. (**C**) Average fluorescence intensity of the SL2-based *dma-1* transcriptional reporter in the PVD cell body in wild type (*dma-1(xr80)*) versus *lin-29(n333lf)* mutants (*dma-1(xr80); lin-29(n333lf)*). Error bars, SEM. NS, not significant by a Student's *t*-test. (**D**) Average dendrite length regrown in the *lin-29(xe61) [lin-29::gfp]* allele without and with the GFP nanobody::ZIF-1 24 hours following dendritomy of the primary dendrite at the young adult stage. Successful *lin-29* knockdown was confirmed by the elimination of LIN-29::GFP proteins from PVD nuclei. Error bars, SEM. NS, not significant by a Student's *t*-test.

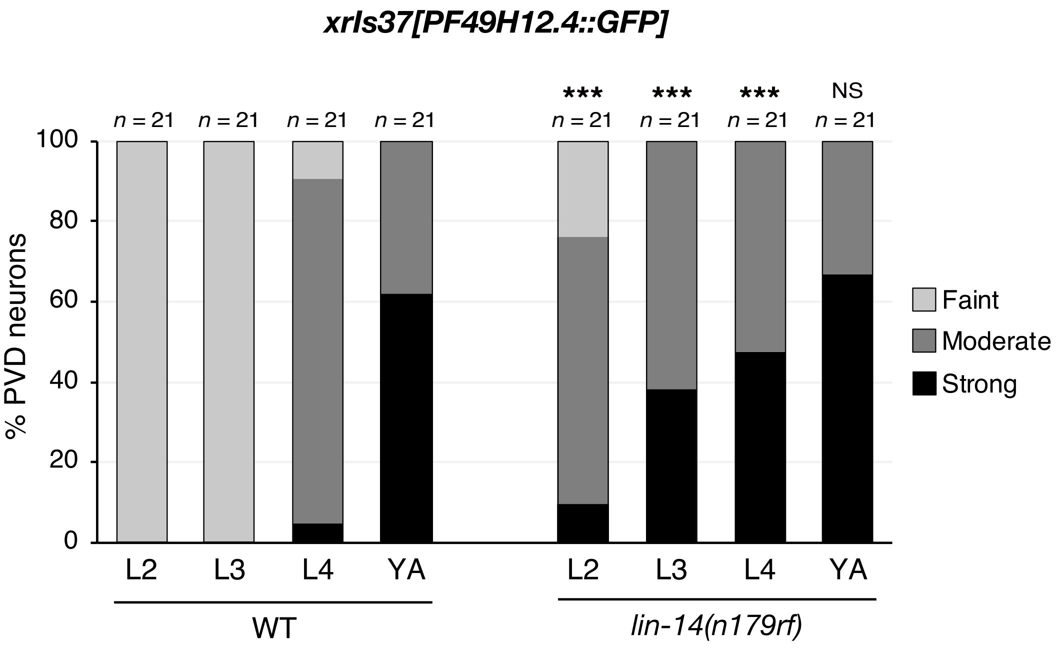


**Figure S6. *lin-14* regulates the timing of PVD neuronal differentiation.** Expression levels of the *xrIs37[PF49H12.4::GFP]* reporter, a PVD maturation marker, in the PVD cell body in wild type and *lin-14(n179rf)* mutants at four different developmental stages. The fluorescence intensity was quantitatively determined to be faint, moderate, or strong. ***$p < 0.001$, relative to wild-type controls at the same stage, by a Student's *t*-test, based on the average fluorescence intensity. NS, not significant.

### **Table S1.** ***C. elegans*** **strains used in this study**

| Strains | Mutations | Integrated transgenes | Extrachromosomal transgenes |
|---|---|---|---|
| XN1588 | | *xrIs21[Plin-4::gfp] X* | *xrEx537[PF49H12.4::mCherry]* |
| XN1540 | | | *xrEx425[Plin-14::gfp]; xrEx537[PF49H12.4::mCherry]* |
| XN1587 | | | *xrEx533[Plet-7::gfp]; xrEx537[PF49H12.4::mCherry]* |
| XN1586 | | | *xrEx117[Plin-41::gfp]; xrEx537[PF49H12.4::mCherry]* |
| XN1427 | | *xrIs21[Plin-4::gfp] X* | *xrEx514[Plin-14::mCherry]* |
| XN1422 | | | *xrEx303[Plet-7::gfp]; xrEx518[Plin-41::mCherry]* |
| XN1808 | *lin-4(e912) II* | *xrIs37[PF49H12.4::gfp] IV* | |
| XN1564 | *lin-14(n355) X* | *xrIs37[PF49H12.4::gfp] IV* | |
| XN1555 | *lin-14(n179) X* | *xrIs37[PF49H12.4::gfp] IV* | |
| XN2033 | *lin-4(e912) II; lin-14(n179) X* | *wyIs581[Pser-2::mCherry]* | |
| XN2044 | | *xrIs37[PF49H12.4::gfp] IV* | *xrEx751[Pser-2::lin-14]* |
| XN2059 | *lin-14(n179) X* | *xrIs37[PF49H12.4::gfp] IV* | *xrEx751[Pser-2::lin-14]* |
| XN1467 | | *xrIs37[PF49H12.4::gfp] IV* | |
| XN1825 | *let-7(n2853) X* | *xrIs37[PF49H12.4::gfp] IV* | |
| XN1547 | *lin-41(n2914)/unc-29(e1072) lin-11(n1281) I* | *xrIs37[PF49H12.4::gfp] IV* | |
| XN2072 | *lin-41(n2914)/unc-29(e1072) lin-11(n1281) I; let-7(n2853) X* | *xrIs37[PF49H12.4::gfp] IV* | |
| XN2818 | | *wgIs535[lin-28 fosmid::ty1::egfp::3x flag]* | *xrEx1157[Pser-2::mCherry]* |
| XN2540 | *lin-41(xr76) [mNG::lin-41] I* | | *xrEx997[Pser-2::mCherry]* |
| XN2637 | *lin-28(n719) lin-41(xr76) I* | | *xrEx1065[Pser-2::mCherry]* |
| XN2690 | *lin-28(n719) I* | | *xrEx1097[Pser-2::gfp]* |
| XN2691 | *lin-28(n719) I* | | *xrEx1098[Pser-2::lin-41; Pser-2::gfp]* |
| XN2693 | *lin-28(n719) I* | | *xrEx1100[Pser-2::lin-41; Pser-2::gfp]* |
| XN2054 | *lin-4(e912) II* | *xrIs37[PF49H12.4::gfp] IV* | *xrEx744[Plet-7::lin-4]* |
| XN2209 | *lin-4(xr70) [Plet-7::lin-4] II* | *xrIs37[PF49H12.4::gfp] IV* | |
| XN2210 | *lin-4(xr71) [Plet-7::lin-4] II* | *xrIs37[PF49H12.4::gfp] IV* | |
| XN2204 | *let-7(xr67) [Plin-4::let-7] X* | *xrIs37[PF49H12.4::gfp] IV* | |
| XN2205 | *let-7(xr68) [Plin-4::let-7] X* | *xrIs37[PF49H12.4::gfp] IV* | |
| XN2670 | *lin-41(xr76) I; lep-5(ny28) X* | | *xrEx997[Pser-2::mCherry]* |
| XN2460 | *lep-5(ny28) X* | *xrIs37[PF49H12.4::gfp] IV* | |
| XN2888 | | | *xrEx1186[Phpo-30::hpo-30::gfp; Pser-2::mCherry]* |
| XN2895 | | | *xrEx1193[Phpo-30::hpo-30::gfp; Pser-2::mCherry]* |
| XN2896 | *lin-14(n179) X* | | *xrEx1186[Phpo-30::hpo-30::gfp; Pser-2::mCherry]* |
| XN2898 | *lin-14(n179) X* | | *xrEx1193[Phpo-30::hpo-30::gfp; Pser-2::mCherry]* |
| XN2899 | *lin-41(n2914)/unc-29(e1072) lin-11(n1281) I* | | *xrEx1186[Phpo-30::hpo-30::gfp; Pser-2::mCherry]* |
| XN2901 | *lin-41(n2914)/unc-29(e1072) lin-11(n1281) I* | | *xrEx1193[Phpo-30::hpo-30::gfp; Pser-2::mCherry]* |
| TV24913 | *dma-1(wy1246) [dma-1::gfp] I* | | |
| XN2870 | *dma-1(wy1246) I; lin-14(n179) X* | | |
| XN2862 | *dma-1(xr80) [dma-1::sl2::mNG] I* | | *xrEx1180[Pser-2::mCherry]* |
| XN2902 | *dma-1(xr80) I; lin-14(n179) X* | | *xrEx1180[Pser-2::mCherry]* |
| PD1301 | *lin-14(cc2841) [lin-14::gfp] X* | | |
| XN1657 | *dma-1(xr50) I* | *xrIs37[PF49H12.4::gfp] IV* | |
| XN2365 | *dma-1(xr50) I; lin-14(n179) X* | *xrIs37[PF49H12.4::gfp] IV* | |
| XN2864 | *dma-1(wy1246) lin-41(n2914)/unc-29(e1072) lin-11(n1281) I* | | |
| XN2920 | *dma-1(xr80) lin-41(ma104) I* | | *xrEx1180[Pser-2::mCherry]* |
| XN1771 | *lin-41(ma104) I* | *xrIs37[PF49H12.4::gfp] IV* | |
| XN2807 | | *xrIs37[PF49H12.4::gfp] IV* | *xrEx1152[Pser-2::dma-1]* |
| XN2810 | | *xrIs37[PF49H12.4::gfp] IV* | *xrEx1155[Pser-2::dma-1]* |
| XN2812 | *lin-41(ma104) I* | *xrIs37[PF49H12.4::gfp] IV* | *xrEx1152[Pser-2::dma-1]* |
| XN2813 | *lin-41(ma104) I* | *xrIs37[PF49H12.4::gfp] IV* | *xrEx1155[Pser-2::dma-1]* |
| XN2725 | *lin-29(xe61) [lin-29 a/b::gfp::3x flag] II* | | *xrEx1126[Pser-2::mCherry]* |
| XN2905 | *dma-1(xr80) I; lin-29(n333) II* | | *xrEx1180[Pser-2::mCherry]* |
| XN2890 | *lin-29(xe61) II* | | *xrEx1188[Pser-2::GFP nanobody::zif-1; Pser-2::mCherry]* |
| XN2892 | *lin-29(xe61) II* | | *xrEx1190[Pser-2::GFP nanobody::zif-1; Pser-2::mCherry]* |

### Table S2. Plasmids used in this study

| Plasmids |
|---|
| *Plin-4::GFP* |
| *Plin-14::GFP* |
| *Plet-7::GFP* |
| *Plin-41::GFP* |
| *PF49H12.4::mCherry* |
| *Plin-14::mCherry* |
| *Plin-41::mCherry* |
| *PF49H12.4::GFP* |
| *Pser-2::lin-14* |
| *Pser-2::mCherry* |
| *Pser-2::GFP* |
| *Pser-2::lin-41* |
| *Plet-7::lin-4* |
| *Plin-4 5' homology arm::Plet-7::SEC::Plin-4 3' homology arm repair template* |
| *Plin-4 sgRNA11 Cas9* |
| *Plin-4 sgRNA12 Cas9* |
| *Plet-7 5' homology arm::Plin-4::SEC::Plet-7 3' homology arm repair template* |
| *Plet-7 sgRNA6 Cas9* |
| *Plet-7 sgRNA8 Cas9* |
| *Phpo-30::hpo-30::gfp* |
| *dma-1 5' homology arm::sl2::mNG::SEC::dma-1 3' homology arm repair template* |
| *dma-1 sgRNA Cas9* |
| *Pser-2::dma-1* |
| *Pser-2::GFP nanobody::zif-1* |
